# Hierarchical control of visual perceptual learning with model-synthesized stimuli

**DOI:** 10.64898/2026.11.23.768712

**Authors:** Younes Valibeigi, Ashim Pandey, Christopher C. Pack, Shahab Bakhtiari

## Abstract

Visual perceptual decisions depend on both how the visual system represents the stimuli and how those representations are read out to guide perception and behavior. Improved visual perception through training, a phenomenon known as Visual Perceptual Learning (VPL), has been partly attributed to adjustments of the readout weights across visual areas to reach better decisions. However, it has been difficult to test whether this reweighting remains at a late stage of the hierarchy or can amplify the readout of whichever hierarchical stage has the most informative representations for the training stimulus. Here, we leveraged DorsalNet, a hierarchical neural network model of the primate dorsal stream, to synthesize motion stimuli that selectively target early or late areas of the primate dorsal pathway. Human observers were trained to discriminate the synthesized stimuli over three days, and transfer was subsequently measured across retinal location and stimulus type. Training improved performance for both stimulus types (early- and late-stage targeting stimuli), but through different mechanisms. Training with late-stage stimuli significantly increased perceptual sensitivity, a gain that transferred across retinal locations. Training with early-stage stimuli produced only a weak, non-significant change in sensitivity; instead, it significantly reduced a conservative response bias, an effect that did not persist at the new retinal location. Specificity indices for stimulus location and type showed different patterns across stimulus categories. A DorsalNet-based model of VPL reproduced both the learning dynamics and these distinct specificity patterns most closely when the model readout was solely plastic for late, rather than early, model layers. Together, these results support a late-stage reweighting account of perceptual learning, in which readout from a preferred, late stage of the dorsal hierarchy is reweighted regardless of the trained stimulus. Under this account, stimuli well matched to the readout stage yield sensitivity gains, whereas poorly matched stimuli afford only weak sensory evidence, prompting observers to compensate by shifting their decision criteria.

## 1 Introduction

Practice on a visual task reliably improves performance, a phenomenon known as visual perceptual learning (VPL) (Ball & Sekuler, 1982; Fiorentini & Berardi, 1980; Lu & Dosher, 2022). These improvements, however, are often narrowly restricted: gains from training on one stimulus or task frequently fail to transfer to closely related stimuli or retinal locations, a property known as specificity (Dosher & Lu, 2020). The degree of specificity reflects how broadly a learned improvement generalizes across the visual system, and it has therefore become a key behavioral tool for inferring where in the cortical hierarchy learning-related plasticity occurs (Lu & Dosher, 2022; Petrov et al., 2005).

A leading account of this plasticity is the reweighting hypothesis. According to this hypothesis, sensory representations and the tuning of their neurons remain largely fixed, and learning instead adjusts the readout weights linking these representations to decision units (Dosher et al., 2013; Lu & Dosher, 2022; Petrov et al., 2005; Sotiropoulos et al., 2011). Reweighting could in principle occur at early or late stages of the visual hierarchy and may involve feedback from higher to lower cortical areas (Dosher et al., 2013). Which stage influences the readout for a given task therefore remains an open question.

The first hypothesis, *adaptive-stage reweighting*, holds that reweighting acts on readouts from all stages of the hierarchy, rather than from a fixed stage, so that the adjusted readout weights come to favor whichever stage is most informative about the training stimulus (Dosher et al., 2013). In motion perceptual learning, this assumption explains observed patterns of specificity well (Bakhtiari et al., 2020), and it agrees with electrophysiological evidence that training can change which areas of the primate dorsal stream contribute to a perceptual decision (Liu & Pack, 2017). Relocating the readout to match each new stimulus requires plasticity distributed across the hierarchy.

The second hypothesis*, late-stage reweighting*, draws on the visual system’s lifelong experience with natural streams of sensory input. Over a lifetime, the brain learns to process this input by relying mainly on higher-order representations in the visual cortex, whether of complex objects in the ventral stream (DiCarlo et al., 2012) or of complex motion patterns such as optic flow in the dorsal stream (Mineault et al., 2012). Due to this lifelong experience and adaptation, the late-stage reweighting hypothesis proposes that perceptual decisions remain anchored to these late-stage representations, regardless of the trained stimulus, and that learning only adjusts how fixed late-stage representations are read out to form decisions. This form of learning parallels the lazy regime described in artificial neural networks (ANNs), in which internal representations remain essentially fixed, and learning is confined largely to the readout weights (Chizat et al., 2019; Farrell et al., 2023; Flesch et al., 2022).

The two hypotheses make diverging predictions for stimuli that are synthesized to maximally engage a single stage of the hierarchy. A stimulus that is maximally discernible by a late-stage visual area should improve sensitivity under both hypotheses, since both predict that learning adjusts the readout at that optimal (late) visual processing stage. However, a stimulus tuned only to be distinguished by an early-stage visual area dissociates the two hypotheses. Adaptive-stage reweighting predicts a sensitivity gain: the readout relocates to the early stage, which is the most informative stage for this stimulus. Late-stage reweighting predicts a smaller or slower sensitivity gain: the readout stays preferentially at the late stage, which is not maximally informative for this stimulus. Conventional stimuli cannot enable this test. Even stimuli designed to probe early stages evoke selective responses at late stages (Adab et al., 2014; Khawaja et al., 2013). Perturbing or inactivating a late-stage area also impairs the discrimination of stimuli designed for an early stage (Adab & Vogels, 2016). Naturalistic stimuli engage many hierarchical levels at once and therefore cannot isolate any single stage.

ANNs offer a way to synthesize the stimuli this test requires. ANNs trained on ecologically relevant tasks have emerged as powerful models of the visual system (Bakhtiari et al., 2021; Khaligh-Razavi & Kriegeskorte, 2014; Mineault et al., 2021; Yamins et al., 2014), capturing key aspects of neural responses and behavior across tasks and species (Geirhos et al., 2021; Schrimpf, Kubilius, Hong, et al., 2020). Their internal layers correspond, at least approximately, to successive stages of the biological visual hierarchy (Mineault et al., 2021; Schrimpf, Kubilius, Hong, et al., 2020; Schrimpf, Kubilius, Lee, et al., 2020; Yamins et al., 2014). Stimulus generation, which optimizes an input to maximally or differentially activate a target unit or layer, can therefore tune a stimulus to a single stage of this hierarchy, or even to a single unit (Bashivan et al., 2019; Olah et al., 2017; Ponce et al., 2019; Walker et al., 2019). Naturalistic stimuli offer no such control, and their engagement of each stage can be measured only where neural recordings are feasible. ANNs have also proven useful for modeling perceptual learning itself, capturing how stimulus variability and task structure shape generalization and specificity in both human observers and model units (Cheng et al., 2025; Cohen & Weinshall, 2017; Liu et al., 2024; Manenti et al., 2023; Shen et al., 2024; Volk et al., 2026; Wenliang & Seitz, 2018). This modeling work, however, has not yet been paired with stimuli synthesized to isolate a single hierarchical stage.

Pairing these approaches requires a visual hierarchy whose stages are well characterized, and the primate dorsal pathway, which processes visual motion, offers such a hierarchy. Motion signals ascend this pathway from primary visual cortex (V1) through the middle temporal area (MT) to the medial superior temporal area (MST) (Maunsell & Essen, 1983). Neurons in V1 respond to the orientation and direction of local features within small receptive fields (Hubel & Wiesel, 1968). Neurons in MT are selective for the direction and speed of motion (Maunsell & Essen, 1983), while neurons in MST integrate motion over wide regions of the visual field (Saito et al., 1986; Tanaka et al., 1986) and respond selectively to optic-flow patterns such as expansion, rotation and translation (Duffy & Wurtz, 1991; Mineault et al., 2012). Receptive fields grow along this progression: MT receptive fields are about ten times larger than V1 receptive fields at the same eccentricity (Albright & Desimone, 1987), and MST receptive fields are larger still (Desimone & Ungerleider, 1986). Learning that draws on a later stage should therefore transfer more broadly across retinal locations. Motion perceptual learning also offers direct evidence for reweighting within this pathway. As monkeys learned to discriminate motion direction, responses changed in the lateral intraparietal area (LIP), which converts motion signals into a choice, but not in MT (Law & Gold, 2008), consistent with learning that changes how MT signals are read out rather than how they are represented.

In this study, we used DorsalNet, an artificial neural network trained to estimate self-motion from naturalistic egocentric videos (Mineault et al., 2021). Its processing stages form a hierarchy of motion processing that was shown to be aligned with motion representation from V1 through MT to MST (Mineault et al., 2021). Using stimulus generation techniques, we synthesized pairs of motion stimuli that evoked maximally distinct activation patterns in either an early layer (res0; V1-like; early-stage) or a late layer (res2; MST-like; late-stage) of the model, so that each pair was maximally discernible only by that layer. We then trained human observers on a same–different discrimination task with one of the two stimulus types, and after training, tested transfer to a new retinal location and to the untrained stimulus type. Under the adaptive-stage reweighting hypothesis, training in either condition should increase sensitivity, and the res0 gains should show greater location specificity, given the smaller receptive fields of earlier stages. Under late-stage reweighting, only res2 training should increase sensitivity, and this gain should transfer across retinal location. Finally, in a computational model, we compared different readout plasticity hypotheses using DorsalNet as the visual backbone and examined which readout architecture best reproduced the pattern of human specificity in our data.

## 2 Method

### Ethics Statement

All procedures involving human observers were approved by the Ethics Committee of McGill University (Research Ethics Board File Number: 21-11-002). Before taking part, every observer gave written informed consent.

### 2.1 Overview

This study combined a psychophysical experiment in human observers with a parallel modeling experiment using an artificial neural network (ANN) model of the primate dorsal visual stream. We first used a stimulus generation method applied to DorsalNet (Mineault et al., 2021) to synthesize motion stimuli tuned to an early (res0) and a late (res2) processing stage of the network (Section 2.2). Human observers were then trained and tested on a same–different discrimination task using stimuli from one of these two categories, with transfer assessed across retinal location and across stimulus category (Section 2.3). In a parallel experiment, we trained binary classifiers reading out from five depths of the same frozen DorsalNet model on an analogous discrimination task, using the same synthesized stimuli, to evaluate which readout depth best reproduces the specificity pattern and learning dynamics observed in human observers (Section2.4). Section 2.5 describes the signal-detection and specificity measures used to analyze both datasets.

### 2.2 Stimulus Synthesis

Detailed implementation, code, and synthesized stimuli are available at https://younesvali.com/projects/dorsalnet/.

#### 2.2.1 Model Architecture and Readout Layers

Motion stimuli were synthesized using DorsalNet (Mineault et al., 2021), a three-dimensional residual neural network trained to estimate an observer’s self-motion parameters from egocentric video streams (Fig 1A). DorsalNet processes video input through an initial convolutional stage (s1), broadly analogous to primary visual cortex (V1), followed by four residual stages (res0–res3). Representational similarity analysis demonstrated alignment between different residual blocks of the trained DorsalNet and V1, MT, and MST areas along the dorsal visual pathway (Fig 1B) (Mineault et al., 2021). In this study, all network weights were frozen throughout stimulus synthesis. For each of res0–res3, we extracted the stage’s activations after its ReLU nonlinearity. Each stage contained 32 channels, and each channel responded across the 11 frames and the spatial (height × width) grid of the input, so every stimulus evoked a 32 × 11 × height × width activation array at each stage. Stimuli were synthesized separately to target res0 ("early-layer" stimuli) and res2 ("late-layer" stimuli).

**Fig 1.**
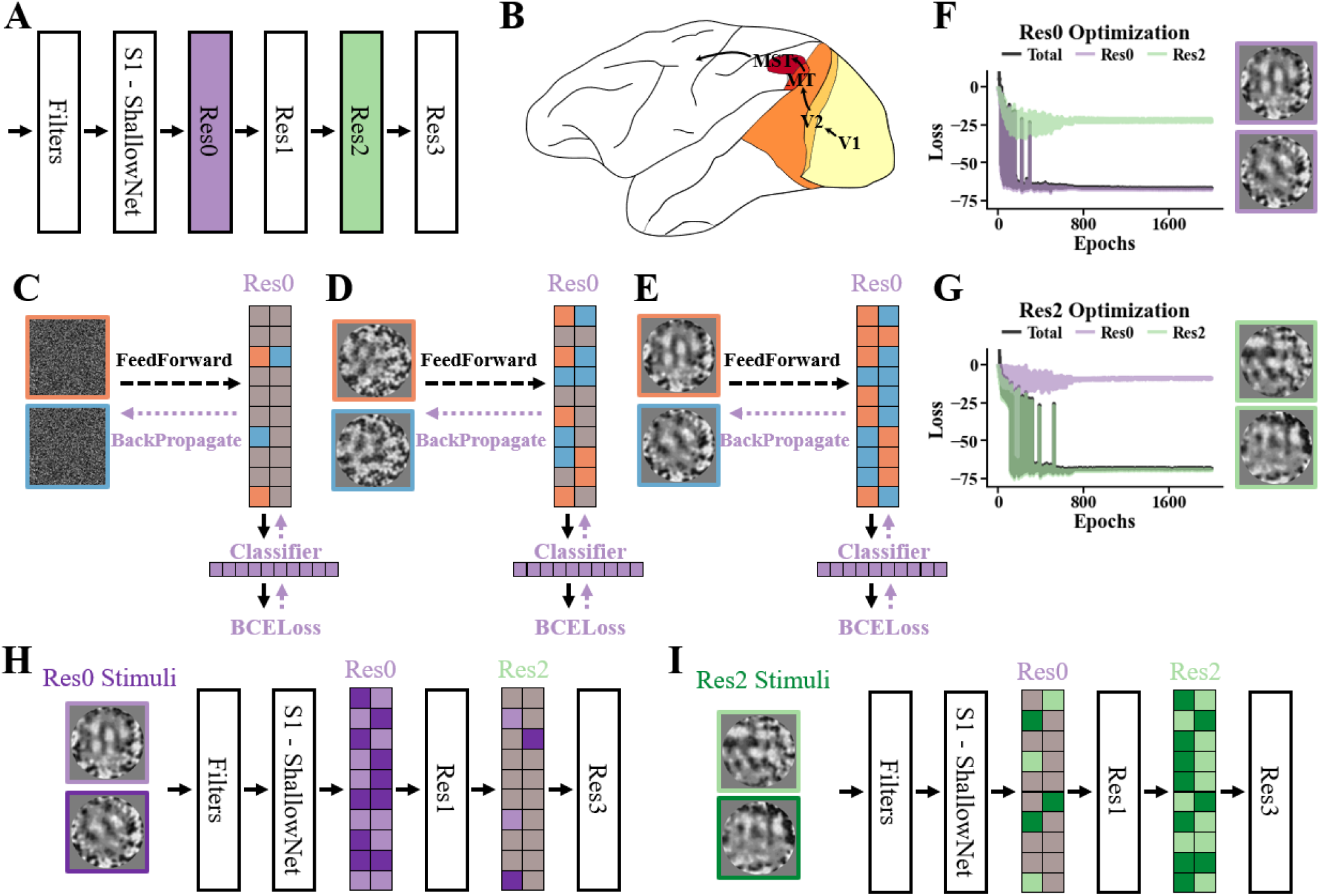
Stimulus synthesis. (A) The DorsalNet model and its hierarchical structure. (B) The dorsal stream in the cortex, which is responsible for motion perception. Representations in areas V1, MT and MST correlate most strongly with those in DorsalNet layers S1, res0 and res1, respectively. (C–E) Stimulus synthesis for layer res0. (C) Two random motion stimuli (orange and blue) are fed to the model. A classifier reads out their res0 activations, and its output enters the BCE loss. The loss is backpropagated to the input, and gradient descent updates the two stimuli while the model weights remain frozen. Initially, the pair does not evoke distinct activations in res0 (mostly gray nodes). (D) As optimization proceeds, the stimuli evoke increasingly distinct res0 activations (more blue and orange nodes). (E) The optimized stimuli evoke distinct activation patterns in the target layer. (F) Loss across optimization iterations for the stimuli targeting res0, shown with the resulting synthesized stimuli. The res0 BCE loss (Section 2.2.3) drove the optimization and was therefore minimized relative to the res2 loss for the same stimulus pair. The total loss, the sum of the res0 BCE loss and the auxiliary penalty terms (Section 2.2.3), closely tracked the res0 loss. (G) Loss across optimization iterations for the stimuli targeting res2, shown with the resulting synthesized stimuli. The res2 BCE loss drove the optimization and was therefore minimized relative to the res0 loss for the same stimulus pair. The total loss closely tracked the res2 loss. (H–I) Schematic of the activations that the synthesized stimuli evoke in res0 and res2. (H) Stimuli optimized for res0 (light and dark purple) evoke maximally distinct activation patterns in res0, but not in res2. The information that distinguishes these stimuli therefore lies in res0, which corresponds to an earlier stage of visual processing. (I) Stimuli optimized for res2 (light and dark green) evoke maximally distinct activation patterns in res2, but not in res0. The information that distinguishes these stimuli therefore lies in res2, which corresponds to a later stage of visual processing.

#### 2.2.2 Linear Readout Classifiers

To generate stimuli targeting a specific processing stage, we first defined, for each of the four residual layers, a fixed linear readout direction in that layer’s activation space (Fig 1C-E). For each layer, we performed principal component analysis (PCA) on the layer’s activations to the videos used to train DorsalNet, which simulate self-motion through virtual environments rendered in a simulator called AirSim (Mineault et al., 2021; Shah et al., 2017). Before the PCA, each 10-frame video was extended to 11 frames by repeating its last frame, and each layer’s activations were averaged over the two spatial dimensions. This averaging collapsed the layer’s spatial (height × width) dimensions while retaining all 32 channels and 11 time frames, yielding a 352-dimensional feature vector (32 channels × 11 frames) for each video. A linear classifier with one output unit and a sigmoid nonlinearity was then defined for each layer, with its weight vector set, without further training, to a single principal component of this decomposition; both the classifier’s weights and its bias term were frozen for the remainder of stimulus synthesis (the classifier itself was never trained, only the stimulus was optimized against it). Several candidate weight vectors (random vectors, all-zero/all-one vectors, and the first several principal components) were compared qualitatively for the clarity and coherence of the resulting synthesized motion; the second principal component (PC2) produced the clearest, most distinct motion and was used for the stimuli reported here.

#### 2.2.3 Optimization Procedure

For each target layer, an initial stimulus batch was constructed from a single random spatiotemporal pattern (3 color channels, 11 frames, 164 × 164 pixels, sampled independently and identically distributed from a standard normal distribution), duplicated to form a pair, one of which was arbitrarily assigned a category label of 0, the other a label of 1. The three color channels matched DorsalNet’s RGB video input, and all stimuli were converted to grayscale before display (Section 2.2.4). Prior to each forward pass, this pair was multiplied by a fixed circular soft-edged spatial mask (sigmoidal fall off, ∼66-pixel radius, centered on the 164 × 164 frame), restricting the region within which pixel structure could be shaped by the optimization, and then subjected to a stochastic random-rescaling transform together with two nested random spatial jitter transforms (integer-pixel translation with wraparound; shifts of up to 6 and 3 pixels, respectively). The maximum jitter magnitude on each iteration was scaled by a logistic decay function of iteration number (steepness = 20, midpoint = 0.25), so that jitter was largest early in optimization and decayed to zero, leaving only the rescaling transform, as optimization proceeded.

On each iteration (Fig 1C-E), the pixel values of the transformed stimulus pair were clipped to [–2, 2] before the pair was passed through the frozen DorsalNet model, and each layer’s activation at the central spatial unit was passed through that layer’s fixed linear classifier. These activations were taken from the single spatial unit at the center of each layer’s activation map, following the single-unit optimization used to probe DorsalNet (Mineault et al., 2021). Each unit responds only to pixels within its receptive field, so reading out the central unit confined the synthesized motion to the center of the frame. For the target layer, a binary cross-entropy-based term drove the two category-labeled stimuli to become maximally separable along the layer’s fixed classifier direction. Four auxiliary penalty terms were applied identically on every iteration: (i) a penalty on the mean-squared pixel value, discouraging deviation from neutral gray; (ii) a penalty on mean per-channel color deviation from neutral, discouraging color casts; (iii) a penalty driving the per-frame contrast of a smoothed version of the stimulus toward a fixed target value; and (iv) an analogous penalty applied to the contrast of a composite stimulus formed by superimposing the two category members (spatially flipped and temporally staggered), driving this composite’s contrast toward a second fixed target value. All terms were combined into a single scalar objective and optimized with Adam (initial learning rate = 0.1), with a step-decay learning-rate schedule (multiplicative decay of 0.5 every 100 iterations, beginning after iteration 1200), for a total of 2000 iterations.

#### 2.2.4 Post-Processing Pipeline

Following optimization, each synthesized stimulus pair was transformed into display-ready motion stimuli through a fixed post-processing pipeline applied identically to all stimuli. Frames (128 × 128 pixels) were first spatially smoothed by convolving each color channel independently (depthwise convolution) twice in succession with a fixed Gaussian low-pass filter (half-width 3 pixels, σ = 1.2 pixels, normalized to unit sum; effective σ ≈ 1.7 pixels), which removed the high-spatial-frequency, pixel-level artifacts that stimulus generation typically introduces. Between the two convolutions, any frame exceeding 128 × 128 pixels was center-cropped back to this size, ensuring that all frames matched the dimensions of the subsequent mask. A circular soft-edged mask (sigmoidal falloff, ∼62-pixel radius, on the 128 × 128 frame) was then multiplied onto each frame, attenuating contrast toward the periphery while leaving the central stimulus region unaffected, so that stimuli had no sharp borders that could serve as an additional cue.

Frames were subsequently upsampled two-fold (128 × 128 → 256 × 256 pixels) via spline interpolation, enlarging the stimuli for presentation without introducing pixelation.. The first and last of the 11 optimized frames were discarded, yielding a 9-frame stimulus: these boundary frames receive incomplete temporal context in DorsalNet’s 3D convolutions and are therefore poorly constrained by the optimization. Each frame was then normalized to a fixed contrast and neutral gray luminance, with two correction passes ensuring that both targets were met. Equating contrast and luminance across stimuli ensured that any difference in learning between res0 and res2 stimuli could not arise from systematic differences in these low-level properties. This final normalization is distinct from, and supersedes, the contrast-related penalty terms used during optimization (Section 2.2.3), which serve only to keep the optimization numerically well behaved rather than to fix the eventual stimulus contrast.

Finally, normalized color frames were converted to grayscale via a standard luminance-weighted transform (0.2989R + 0.5870G + 0.1140B), which removed color as a potential cue so that observers had to discriminate the stimuli on the basis of their motion. Grayscale frames were then remapped for display (pixel intensity x → 0.5 + 0.25x, clipped to [0, 1]) and saved as 9-frame GIF animations.

#### 2.2.5 Dissimilarity-Level Generation

To enable observer performance to span a broad dynamic range throughout training, we generated a graded series of stimulus pairs with progressively increasing similarity to the fully optimized target pair for each layer (Fig 2A). As this similarity to the target pair increased, the dissimilarity between the two members of each pair increased as well, and we refer to this graded dimension as the dissimilarity level. To do so, we initialized a second pair of stimuli using the same procedure described above (two identical copies of a single random spatiotemporal pattern, 164 × 164 pixels, 11 frames), together with the same circular mask and Gaussian filter. The activations of the two members of the already-optimized target pair (Section 2.2.3) were computed once, over the target layer’s full activation map (all channels, frames and spatial positions), and held fixed thereafter. The new pair was then optimized, again with Adam optimizer (Kingma & Ba, 2014), a gradient-descent algorithm that adapts the step size of each parameter during optimization (learning rate = 0.001), such that each of its two members independently minimized the Euclidean (L2) distance between its own activation and the corresponding member of the fixed target pair (i.e., the first new stimulus was driven toward the first target stimulus’s activation, and the second toward the second target’s activation) subject to the same four auxiliary penalty terms described in Section

**Fig 2.**
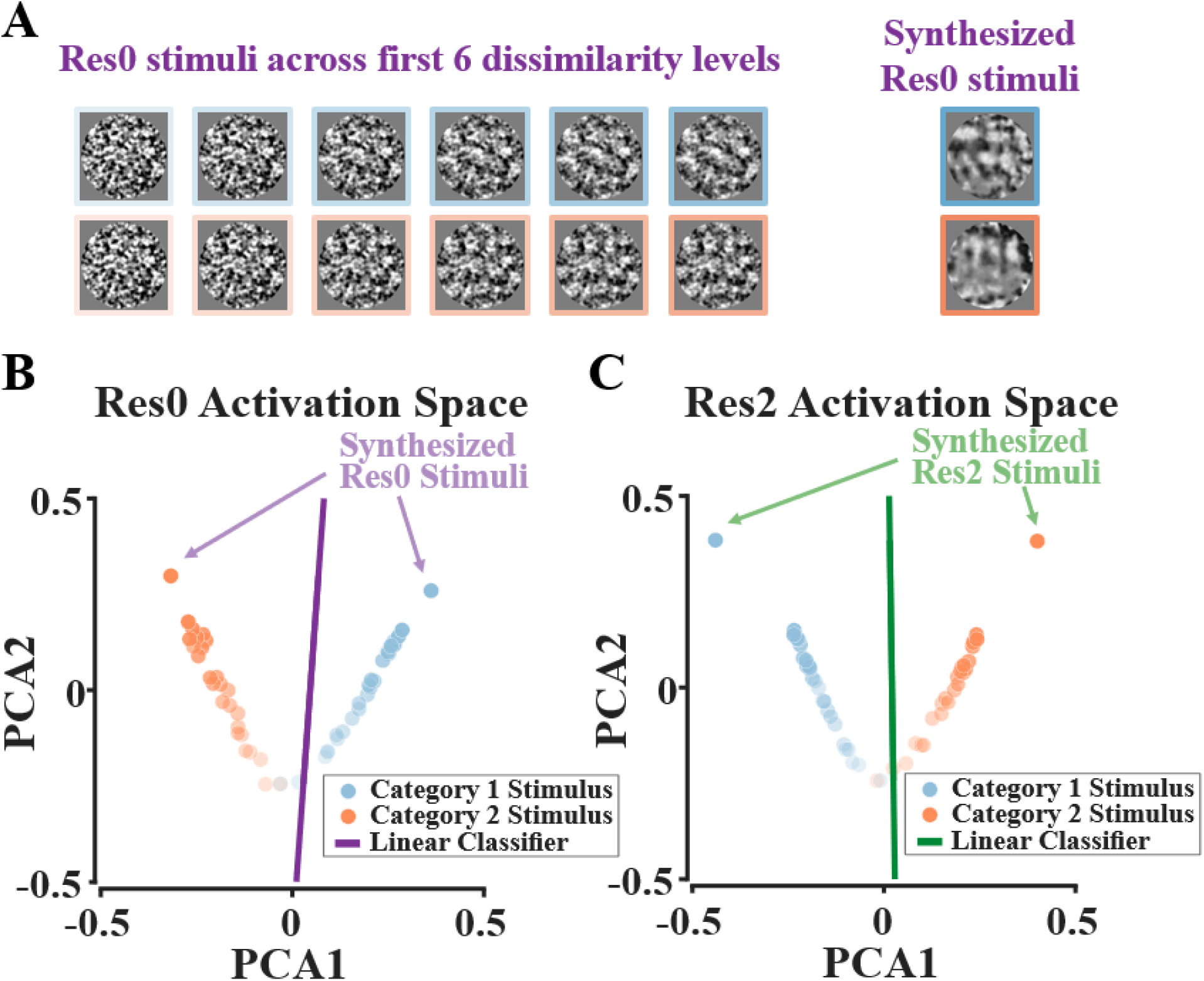
Dissimilarity-level generation. (A) Example set of res0 dissimilarity levels used in the human experiments. For each target layer, the human experiments used the first six generated dissimilarity levels. (B) Activation space of res0 during dissimilarity-level generation for the res0 category. The fully synthesized pair was generated to be as distinct as possible according to the linear classifier. The dissimilarity levels were generated by progressively optimizing from a random initialization toward this pair, yielding a graded sequence of decreasing difficulty (greater transparency indicates a lower dissimilarity level). (C) Activation space of res2 during dissimilarity-level generation for the res2 category.

#### 2.2.3. The stochastic jitter and rescaling transforms were carried over at the fixed magnitude reached at the end of the preceding optimization stage (Section 2.2.3), rather than following a new decay schedule

The evolving pair was saved every 50 iterations, producing a sequence of paired stimuli of progressively decreasing activation distance from the initial random pattern toward the fully optimized target pair (Fig 2B-C) (i.e. from highly noisy, difficult-to-discriminate stimuli to increasingly target-like, easier-to-discriminate stimuli), though this distance was optimized jointly with the same four auxiliary penalty terms described above. This procedure was applied separately for the res0- and res2-tuned target pairs, and the first six saved pairs from each sequence were used in the human experiments described below.

### 2.3 Psychophysical Experiment

#### 2.3.1 Observers

Nineteen observers (undergraduate and graduate students) with normal or corrected-to-normal vision took part (12 males, 7 females). All observers gave written informed consent before participation. The study was approved by the Ethics Committee of McGill University (Research Ethics Board File Number: 21-11-002).

#### 2.3.2 Apparatus

The experiment was conducted in person and controlled with PsychoPy (2024). Eye position was monitored with an infrared eye-tracking system (EyeLink 1000, SR Research; 1,000 Hz sampling rate). Observers viewed the display from 57 cm on a 28-inch monitor (24.6 × 13.7 inches), refresh rate 60 Hz.

#### 2.3.3 Stimuli

Stimuli subtended 5° × 5° of visual angle at the 57 cm viewing distance, presented against a uniform gray background. During training and during the stimulus-category transfer test, stimuli appeared in the upper-right visual field, at a reference angle of 45° and an eccentricity of [+5°, +5°] from screen center. During the location-transfer test, stimuli instead appeared in the lower-right visual field, at a reference angle of 315° and an eccentricity of [+5°, –5°].

#### 2.3.4 Procedure

##### 2.3.4.1 Training

Observers performed a same-different task (Fig 3A). On each trial, observers fixated a black-and-white bullseye at the center of the screen for 1000 ± 200 ms. A stimulus was shown for 150 ms, followed by an inter-stimulus interval of 1000 ± 200 ms, and then a second stimulus for 150 ms. After a further 750 ± 200 ms, observers reported via keypress whether the two stimuli were the same ("s") or different ("d"). During training, correct responses were followed by a brief green ring around fixation with an accompanying tone, and errors by a red ring; this feedback was omitted during the transfer sessions on the fourth day. The inter-trial interval was 750 ms.

**Fig 3.**
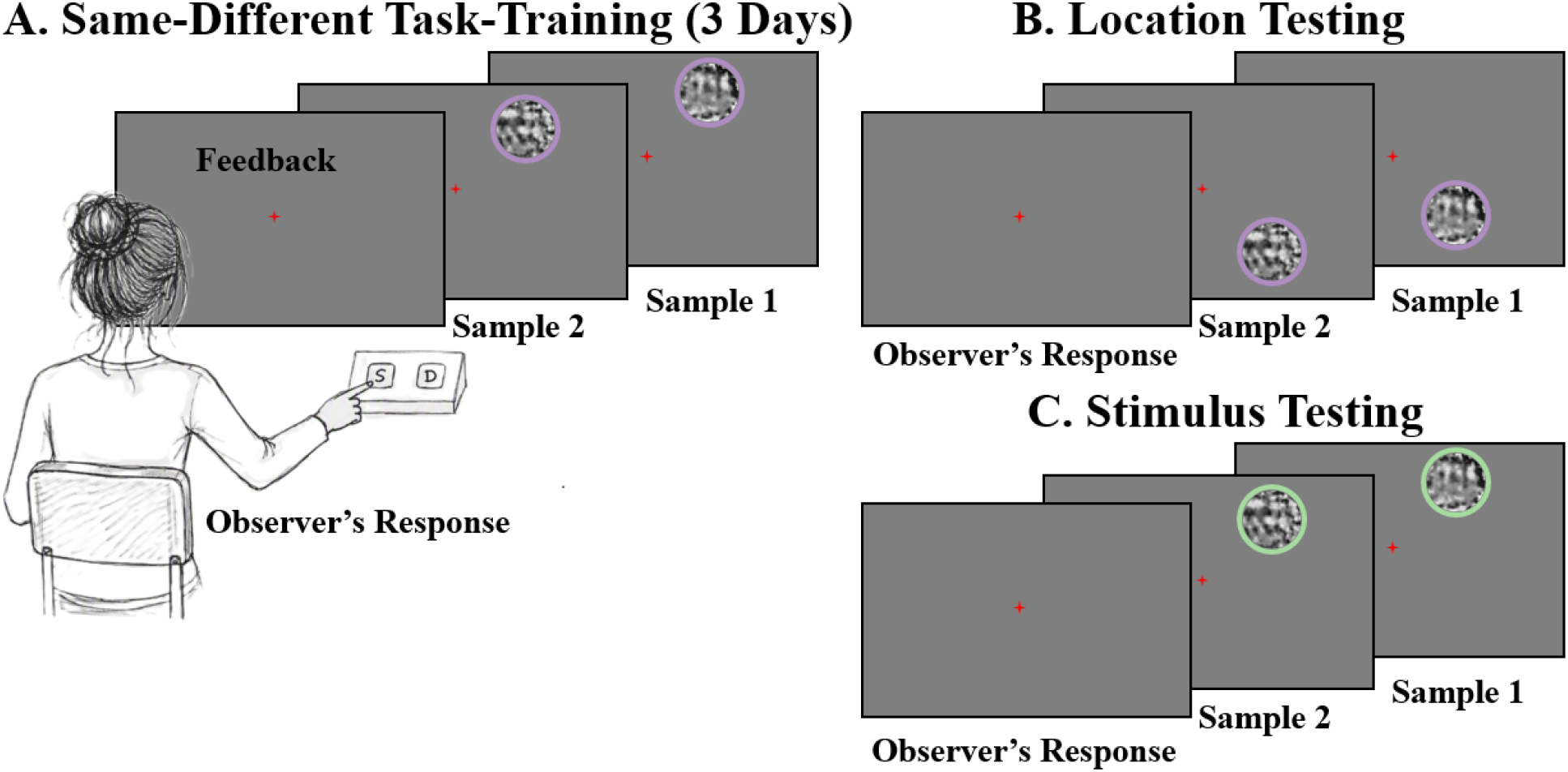
Schematic of same-different task. (A) Training phase (3 days): on each trial, the subject reported whether two sequentially shown video samples belonged to the same category or different categories, with trial-by-trial feedback provided. (B) Location testing: on the fourth day, subjects performed the same task with the stimulus location moved, with no feedback provided. (C) Stimulus testing: subjects performed the task with the stimulus category switched relative to training (i.e., res0-trained subjects tested on res2 stimuli and vice versa), with no feedback provided.

Observers were randomly assigned to one of two stimulus categories (res0 or res2), with 9 observers assigned to the res0 condition and 10 to the res2 condition. Each observer completed one training session per day on three consecutive days, comprising three sets of three 24-trial blocks (216 trials total, 72 per set), with a 1-minute rest between sets. Each block consisted of the six stimulus pairs, with each pair presented in the four possible trial combinations (AA, AB, BA, and BB), yielding 24 trials per block.

##### 2.3.4.2 Testing

On the fourth day, observers completed two transfer sessions, each with the same block/set structure as training (three sets of three 24-trial blocks). In the first session (Fig 3B), location transfer was assessed: the trained stimulus category was presented at the transfer location described in Section 2.3.3. In the second session (Fig 3C), stimulus-category transfer was assessed: stimuli were presented at the original training location, but the untrained category (res0 ↔ res2) was substituted for the trained one.

### 2.4 ANN Readout Modeling Experiment

#### 2.4.1 Readout Architecture and Conditions

We asked whether a classifier reading out from the same frozen DorsalNet model could reproduce the specificity pattern observed in human observers, and how this depended on the readout architecture. For each stimulus category (res0-trained, res2-trained), we compared six readout architectures that differed only in which layer(s) of DorsalNet’s frozen activations they read out from. Five of these architectures each used a single classifier: four read out from one residual layer (res0, res1, res2 or res3), and the fifth read out from all four residual layers concatenated (“All”). The sixth, stimulus-driven architecture used two classifiers, one reading out from res0 and one from res2. Each of these classifiers was paired with the stimuli synthesized for its own layer, so that res0 stimuli were read out from res0 and res2 stimuli from res2.

Each classifier consisted of a single linear layer with a sigmoid output. Its input was the activation at the central spatial unit of the relevant layer(s), following the same central-unit readout used to define the stimulus-synthesis classifiers (Section 2.2.2). Reading out this single unit removed the layer’s spatial (height × width) dimensions while retaining all 32 channels and 11 time frames, yielding 352 input features for a single-layer condition and 1,408 for the four-layer “All” condition. The classifiers’ weights were the only trained parameters here; DorsalNet’s own weights remained frozen throughout. These weights were initialized to zero. Initialization does not alter the final performance of such single-layer linear classifiers, but a common zero starting point ensured that any differences between readout layers early in training arose from the layers’ activations rather than from the initial weights. We restricted the readout-architecture comparison to the four residual layers (res0–res3) and excluded the earlier *s1* layer, whose activations have a substantially larger number of channels, as well as a different spatial resolution, than the residual layers; collapsing across these dimensions at the central spatial unit would yield a much larger feature vector for *s1* than for any res layer, complicating direct comparison across layers.

#### 2.4.2 Training Procedure

Each classifier performed binary classification of individual stimuli into their category of origin. Input stimuli matched the spatial format used in stimulus generation (Section 2.2.3): 164 × 164 pixels, with the stimulus itself confined to a central circular region of 66-pixel radius. The training set for a given layer condition comprised the fully optimized target pair (Section 2.2.3), each stimulus labeled according to which member of the original target pair (label 0 or label 1) its generative trajectory approached. On each training step, the classifier received both stimuli of the training pair, and its weights were updated after every step. The stochastic random-rescaling transform (Section 2.2.3) was applied to each stimulus, and the batch was passed through the frozen model and classifier to obtain predicted category probabilities, which were compared to the true labels using a binary cross-entropy loss. Classifier weights were updated via Adam (learning rate = 0.00001) for 10000 epochs. On every training and evaluation pass, a circular patch of grayscale Gaussian noise (SD = 0.01, radius = 66) was added to the stimulus.

#### 2.4.3 Transfer Testing

After training, each classifier was evaluated, without further weight updates, on three sets mirroring the human transfer design: (i) the training set itself, establishing end-of-training ("learning") performance; (ii) a location-transfer set, in which the same category’s stimuli were shifted from the central position used in training to the bottom-right of the 164 × 164 input grid, mimicking the location-transfer manipulation used with human observers; and (iii) a stimulus-category-transfer set, consisting of the other category’s full stimulus set (e.g., the res2-trained classifier was tested on res0 stimuli, and vice versa) at the original training location. On each evaluation, we recorded a loss measure (mean absolute difference between predicted probability and true label), which was used to compute the specificity index (Section 2.5.2).

Unlike the human transfer stimuli, which occupied a fully distinct region of the visual field, the shifted ANN stimuli partially overlapped the training location: with a 66-pixel radius, each stimulus spanned most of the 164-pixel input grid. This overlap was identical across readout conditions and therefore does not affect the comparison across readout depths.

### 2.5 Data Analysis

#### 2.5.1 Signal Detection Measures

Same–different performance was analyzed using the independent-observation signal detection model (Hautus et al., 2021) (pp. 207–211). Trials on which the two stimuli differed in category (⟨S1,S2⟩ or ⟨S2,S1⟩) were treated as signal-present ("different") trials, and trials on which the two stimuli matched (⟨S1,S1⟩ or ⟨S2,S2⟩) as signal-absent ("same") trials. Hit and false-alarm rates were computed as:

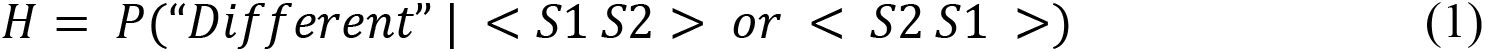

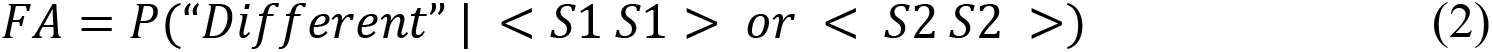

(Hautus et al., 2021) (Eq. 9.1, p. 207). Sensitivity (d′) was estimated via the maximum-proportion-correct transform of H and FA (Hautus et al., 2021) (Eq. 9.3, pp. 208–211):

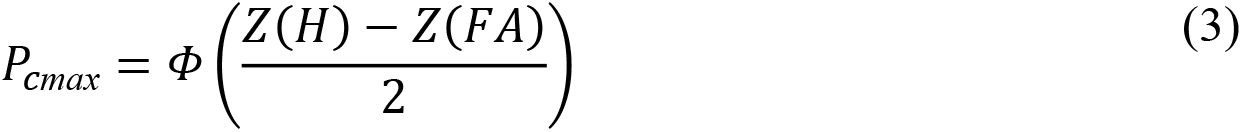

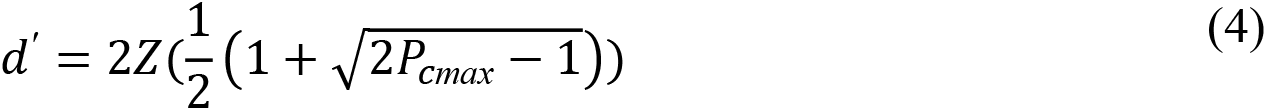

where Φ(·) is the cumulative normal distribution. Response criterion, quantifying response bias independently of sensitivity, was computed as:

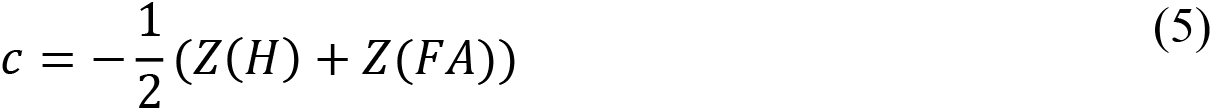

(Hautus et al., 2021) (Eq. 2.1). Under this model, observers are assumed to form independent internal representations of each stimulus, classify each representation into one of the two categories, and respond “different” when the two classifications disagree. The alternative differencing model instead assumes that the observer subtracts the two internal observations and responds “different” when the magnitude of this difference exceeds a criterion (Hautus et al., 2021). Classifying each stimulus matches the assumption that underlies this study: a decision unit reads out a category from the sensory representation of each stimulus (Dosher & Lu, 1999; Petrov et al., 2005). We therefore used the independent-observation model. Under this model, *d′* reflects the perceptual separability of the two categories independent of response bias, whereas *c* reflects the observer’s tendency to favor “same” or “different” responses independent of sensitivity.

#### 2.5.2 Specificity Index

Observers showed an overall bias toward "same" responses throughout the experiment. Performance measures based on same-category trials are therefore confounded by this bias: a "same"-biased observer would score well on same trials through response tendency alone, independent of genuine discrimination ability. To isolate discrimination performance from this criterion effect, we restricted the specificity index below to different-category trials only, for the human data. For human observers, the relevant performance measure was the hit rate H (Eq. 1), computed at the training or transfer stage of interest. For the ANN classifiers, whose task is a direct single-stimulus classification (Section 2.4.1) rather than a same–different judgment, no analogous same-response bias applies; here, the corresponding measure was instead computed as one minus the mean absolute prediction error across the full relevant evaluation set.

The specificity index was defined as:

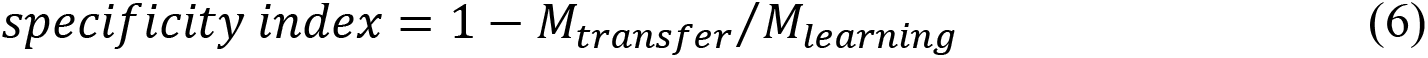

(Lu & Dosher, 2022), where M_learning_ is this performance measure at the end of training, and M_transfer_ is the same measure computed on the relevant transfer set (location or stimulus-category) (Fig 4B). This index was computed separately for location-transfer and stimulus-category-transfer, for both the res0- and res2-trained conditions (human data) and for all five readout-depth conditions (ANN data).

**Fig 4.**
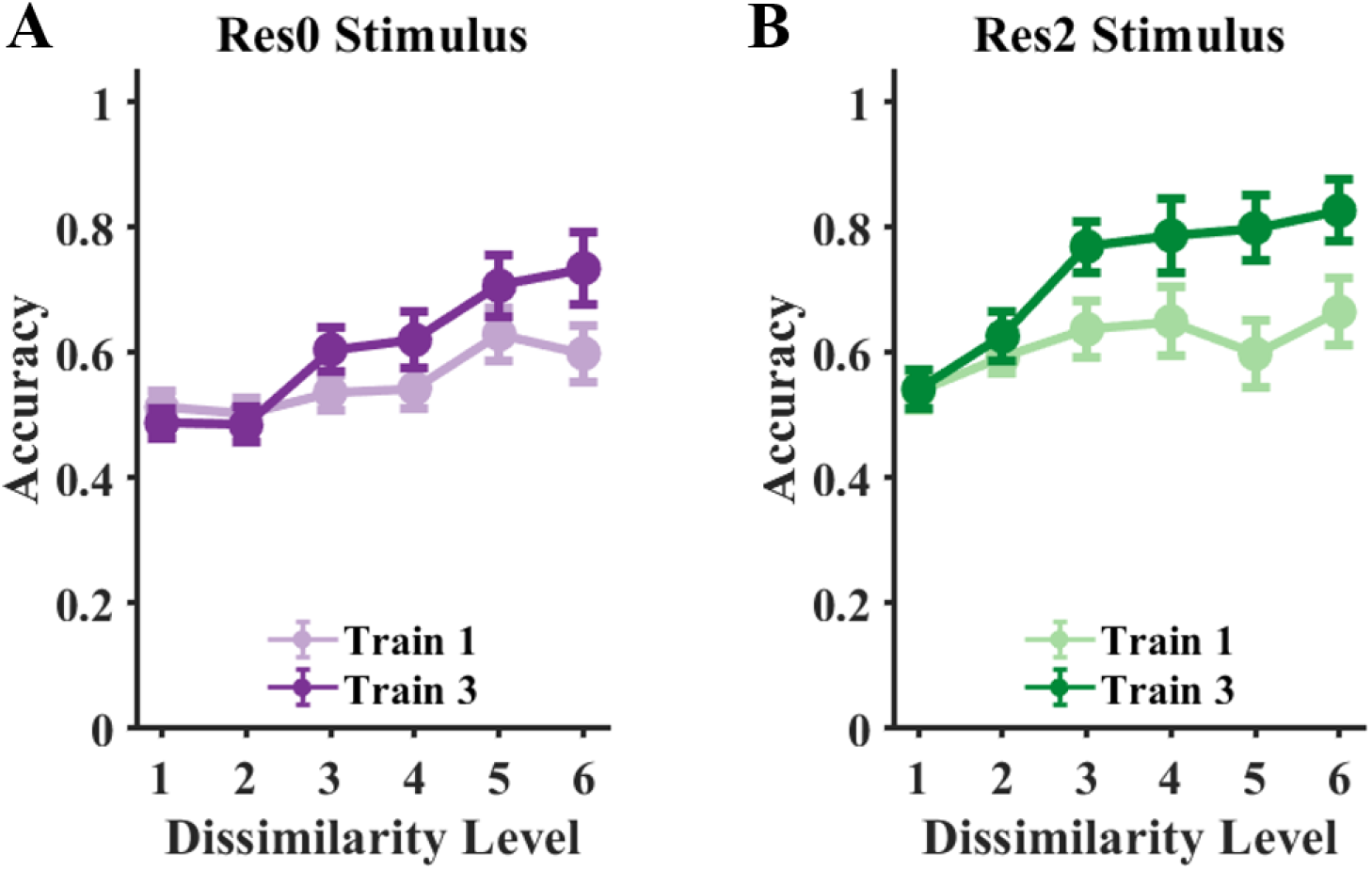
Psychometric functions across training. (A) Performance across six dissimilarity levels for subjects trained on res0 stimuli, before (Train 1) and after (Train 3) training. Performance improved after training, particularly for levels 3–6. (B) Performance across six dissimilarity levels for subjects trained on res2 stimuli. As for res0, performance improved after training, particularly for levels 3–6.

#### 2.5.3 Statistical Analysis

Sensitivity (d′) and criterion (c), computed under the independent-observation model (Section 2.5.1), were each analyzed using paired-samples t-tests, separately for the res0 and res2 stimulus categories, to test for change across three within-subject comparisons: Train 1 vs. Train 3 (learning during training), Train 3 vs. Location Test (transfer across retinal location), and Train 3 vs. Stimulus Test (transfer across stimulus category). Sensitivity was additionally compared between the res0 and res2 stimulus categories, at the end of training (Train 3) and at location testing, using independent-samples t-tests. All t-tests were two-tailed, with degrees of freedom determined by the number of participants providing valid paired (or, for between-group comparisons, unpaired) observations for each specific test.

The qualitative correspondence between human and ANN specificity-index orderings across readout depths (Section 3.6) was assessed by directly comparing the pattern of indices across conditions, rather than via a formal statistical model-comparison procedure.

## 3 Results

To determine at which stage of the visual hierarchy readout reweighting operates during VPL, we used stimulus generation with DorsalNet (Mineault et al., 2021), a neural network model of the primate dorsal stream. For each of two layers, an early layer (res0) and a late layer (res2), we synthesized a pair of dynamic stimuli that evoked maximally distinct activation patterns only in that layer. We asked whether, during training, the neural process of learning could find the optimal stage of the hierarchy and read out from that stage to improve perceptual performance. Human observers were trained on a same–different discrimination task with one of the two stimulus categories (res0 or res2) and were then tested for transfer to a new retinal location and to the untrained stimulus condition. The transfer tests gave us diagnostic tools to study the readout plasticity during learning, as we will explain in the following sections.

### 3.1 Stimulus Optimization

The representations in DorsalNet’s layers align hierarchically with areas of the primate dorsal stream (Mineault et al., 2021). We used this model to synthesize motion stimulus pairs that evoked maximally distinct activation patterns only in an early (res0) or a late (res2) layer. For each targeted layer, a stimulus pair (two multi-frame videos) were optimized to produce maximally distinct activation patterns at that layer, following the procedure detailed in Sections 2.2.2–2.2.4. In both optimizations, the classification loss fell further at the target layer than at the non-target layer (Fig 1F, G). This target-layer loss also closely tracked the total loss, which additionally included the auxiliary penalty terms (Section 2.2.3).

To capture a broad range of observer performance, we synthesized a graded series of stimulus pairs for each target layer of the model, ranging from near-identical noisy pairs to the fully optimized pair (Section 2.2.5)(examples at https://younesvali.com/projects/dorsalnet/). The human experiments used the first six dissimilarity levels of each series; Fig 2A shows these levels for res0, alongside the fully optimized pair. To visualize how the target layer distinguished these levels, we performed Principal Component Analysis (PCA) on that layer’s activations elicited by the generated stimuli and projected both the stimuli activations and the fixed linear classifier (Section 2.2.2) into the first two principal components. In these projections (Fig 2B and C for res0 and res2, respectively), the dissimilarity levels formed a graded sequence, from noisy pairs near the classifier boundary to the widely separated, fully optimized pair.

### 3.2 Performance improves with training for both stimulus categories

Observers performed a same-different task over three consecutive days (Train 1–Train 3). On each trial, observers judged whether two sequentially presented motion stimuli were the same or different. Stimulus pairs were drawn from six graded dissimilarity levels (Fig 2A), ranging from near-identical to maximally distinct. Trial-by-trial feedback was provided during training sessions but withheld during transfer testing. Fig 4 shows observers’ performance across the six dissimilarity levels for both the res0 and res2 categories. Performance remained near chance at the most difficult levels, but by the third day of training, observers had learned to discriminate the stimulus pairs, and performance increased markedly for levels 3–6 in both categories.

### 3.3 Performance gains arise from increased sensitivity for res2 but from reduced response bias for res0

Performance in a same–different task depends on two quantities: sensitivity (d′), which reflects the perceptual separability of the two stimuli in a pair, and response criterion (c), which reflects the observer’s tendency to favor "same" or "different" responses (Section 2.5.1). Adjusting both sensitivity and response criterion can lead to improved performance (Laamerad et al., 2024; Laamerad et al., 2025). To determine which of these quantities produced the performance gains in each stimulus category, we estimated both with the independent-observation signal detection model (Hautus et al., 2021) (Section 2.5.1), separately for res0- and res2-trained observers. Training with res2 stimuli increased sensitivity from Train 1 to Train 3 (Train 1 vs. Train 3, paired-samples t-test, *t*(9) = 4.50, *p* = .001), whereas training with res0 stimuli produced only a weak, non-significant change (Train 1 vs. Train 3, paired-samples t-test, *t*(8) = 1.86, *p* = .100) (Fig 5A). Across all three training sessions, res2-trained observers showed numerically higher sensitivity than res0-trained observers. This difference was already apparent, though smaller, at Train 1 (*d*^′^ = 1.05 vs. = 0.53), although it was not statistically significant (res2 vs. res0, independent-samples t-test, *t*(17) = 1.56, *p* = 0.136, *n* = 9 (*res*0), *n* = 10 (*res*2))) and is consistent with a default late-stage readout: stimuli synthesized to be maximally discernible at a late stage should be more discriminable from the outset (Section 4.1). By the end of training (Train 3), discrimination was significantly better for res-2 stimuli (res2 vs. res0, independent-samples t-test, *t*(17) = 2.41, *p* = 0.028, *n* = 9 (*res*0), *n* = 10 (*res*2)), consistent with perceptual decisions in this task being read out from a late rather than an early stage.

**Fig 5.**
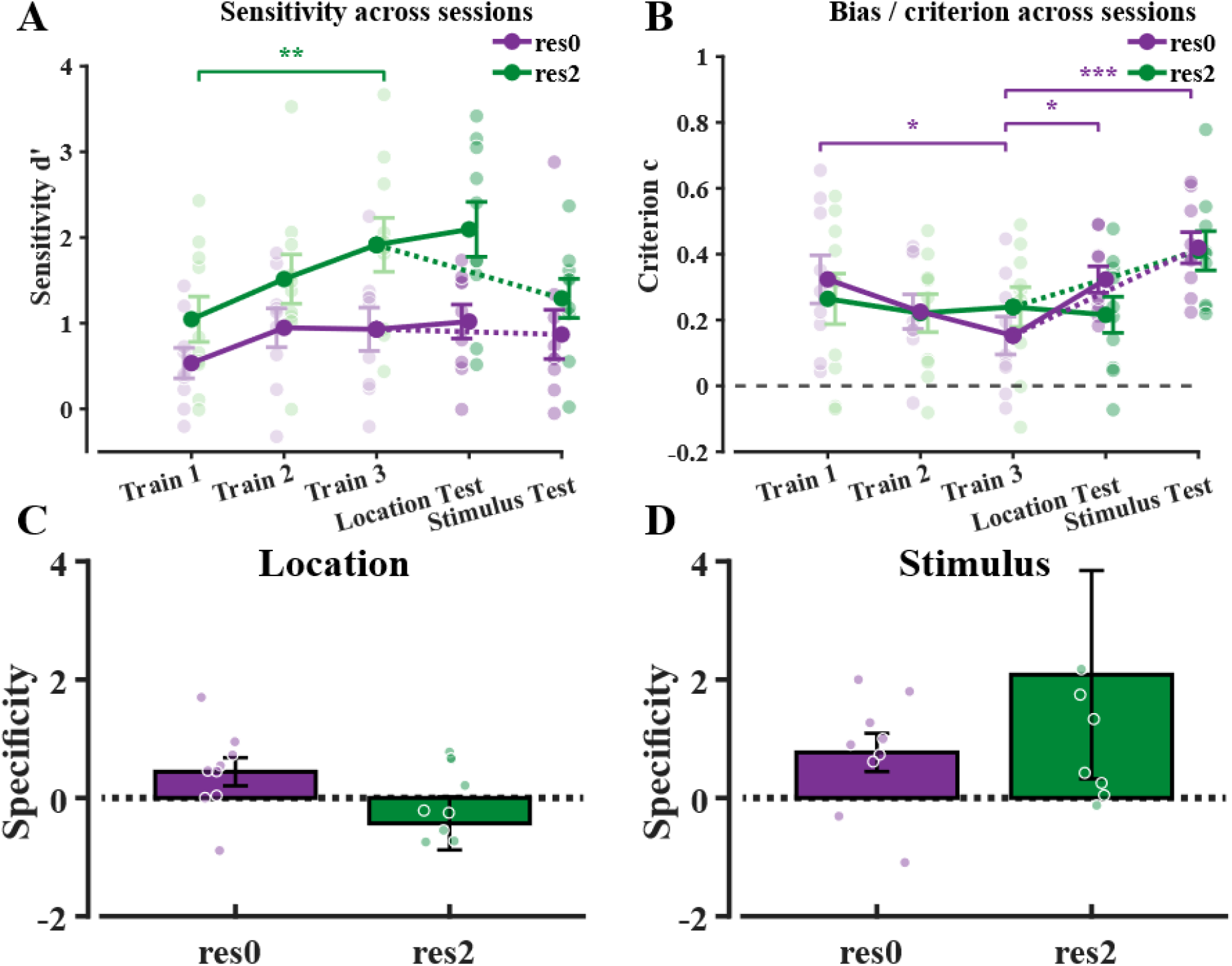
Psychophysics results from the human same–different task, computed using the independent-observation signal detection model. (significance indicated as * p < .05, ** p < .01, *** p < .001). (A) Sensitivity (d′) across sessions (Train 1–3, Location Test, Stimulus Test) for res0- and res2-trained observers. Res2 sensitivity was higher than res0 sensitivity in every session, and this difference reached significance at the end of training (Train 3) and at location testing. Res2 sensitivity increased significantly across training (Train 1 vs. Train 3, **) and was maintained at location testing without further significant change. Res0 sensitivity did not change significantly across training or transfer. (B) Response criterion (bias) across the same sessions. Observers in both categories were initially biased toward "same" responses. For res0, this bias decreased significantly across training (*) and reverted significantly toward baseline at location testing (*). For res2, bias changed little in either comparison. Switching stimulus category increased bias significantly for res0 (***) and marginally for res2. (C, D) Specificity indices, computed from different trials only (Section 2.5.2), for location transfer (C) and stimulus-category transfer (D). The two transfer types showed opposite orderings: specificity was higher for res0 in location transfer but higher for res2 in stimulus-category transfer. In panels C and D, a small number of individual data points falling well outside the plotted axis range were omitted for visual clarity; all statistical analyses used the complete dataset.

Response criterion showed the opposite pattern (Fig 5B). At the start of training, observers in both categories were biased toward reporting "same," indicating a conservative criterion: they required strong evidence before reporting “different.” With training, this bias declined steadily for res0-trained observers (Train 1 vs. Train 3, paired-samples t-test, *t*(8) = 2.99, *p* = .017) but changed little for res2-trained observers (Train 1 vs. Train 3, *t*(9) = 0.45, *p* = .664). Performance therefore improved in both categories, but through different mechanisms: res2 training increased sensitivity, whereas res0 training reduced response bias.

### 3.4 Only the sensitivity gain from res2 training persists on the transfer day

The two mechanisms also differed in their transfer to a new retinal location. After the location change on the transfer day, res2 sensitivity remained elevated and increased numerically relative to the end of training, despite the removal of trial-by-trial feedback (Fig 5A). This pattern is consistent with evidence (Petrov et al., 2006; Seitz & Watanabe, 2003) that perceptual learning can occur without external feedback. For testing at the transfer location, res2 sensitivity was significantly higher than res0 sensitivity (independent-samples t-test, *t*(17) = 2.78, *p* = 0.013, *n* = 9 (*res*0), *n* = 10 (*res*2)), indicating that the res2 sensitivity gain was not tied to the trained retinal location. Res0 sensitivity, in contrast, remained at its low end-of-training level, neither improving nor declining; res0 training had produced little sensitivity gain to begin with, leaving little for a location change to disrupt.

The res0 bias reduction, however, did not survive the transfer test, in which both retinal location and feedback changed (Petrov et al., 2006): bias increased toward its pre-training level (Train 3 vs. Location Testing, *t*(8) = −2.74, *p* = .026), whereas res2 bias remained stable (Train 3 vs. Location Testing, *t*(9) = 0.33, *p* = .750) (Fig 5B). Of the two forms of improvement, only the sensitivity gain from res2 training persisted on the transfer day.

### 3.5 Transfer across stimulus category is asymmetric for sensitivity

Transfer to the untrained stimulus category was asymmetric (Fig 5A). In observers trained on res2, after switching from res2 to res0, sensitivity declined numerically, although this decline did not reach significance (Train 3 vs. Stimulus Test, paired-samples t-test, *t*(8) = 1.26, *p* = .243). In observers trained on res0, switching from res0 to res2, sensitivity did not change (*t*(8) = 0.22, *p* = .830) (Fig 5A). Response bias, in contrast, increased after switching in both directions (Fig 5B), and this increase was significant for res0-trained observers (*t*(8) = −9.88, *p* < .001) and marginal for res2-trained observers (*t*(8) = −2.24, *p* = .055). To compare these patterns with the location-transfer results, we computed a specificity index (Eq. 6) for each transfer type and training category, using only different trials (Section 2.5.2). Location specificity was higher for res0 than for res2, whereas stimulus-category specificity was higher for res2 than for res0 (Fig 5C).

Although these two orderings might appear contradictory, both are consistent with a single account in which perceptual decisions are read out preferentially from a late stage of the visual hierarchy (Section 4.3).

### 3.6 The human specificity pattern is best reproduced by readouts from deep ANN layers

Under the assumption that perceptual learning preferentially reweights higher-level readouts, an ANN readout with access to deeper layers should better reproduce the specificity profile observed in the human data. Under adaptive-stage reweighting, by contrast, a readout from the layer each stimulus was optimized for should better reproduce it. To test these predictions, we trained binary classifiers that read out from individual residual layers of DorsalNet (res0–res3), from all four layers combined, or from the stimulus-driven architecture, which reads out from res0 for res0 stimuli and from res2 for res2 stimuli (Fig 6A). Each architecture implements a different hypothesis about where reweighting acts. Readouts from res2 or res3 implement late-stage reweighting, and readouts from res0 or res1 its early-stage counterpart. The stimulus-driven architecture implements adaptive-stage reweighting: its readout follows whichever layer each stimulus was optimized for. The all-layer readout is the most flexible of the six, since learning determines which layer, or combination of layers, the decision draws on. All classifiers were trained on the fully optimized res0 and res2 target pairs, with weights updated after every training step. For each readout model, we computed the same specificity index for simulated location and stimulus-category transfer as for the human data (Fig 6B, Section 2.5.2).

**Fig 6.**
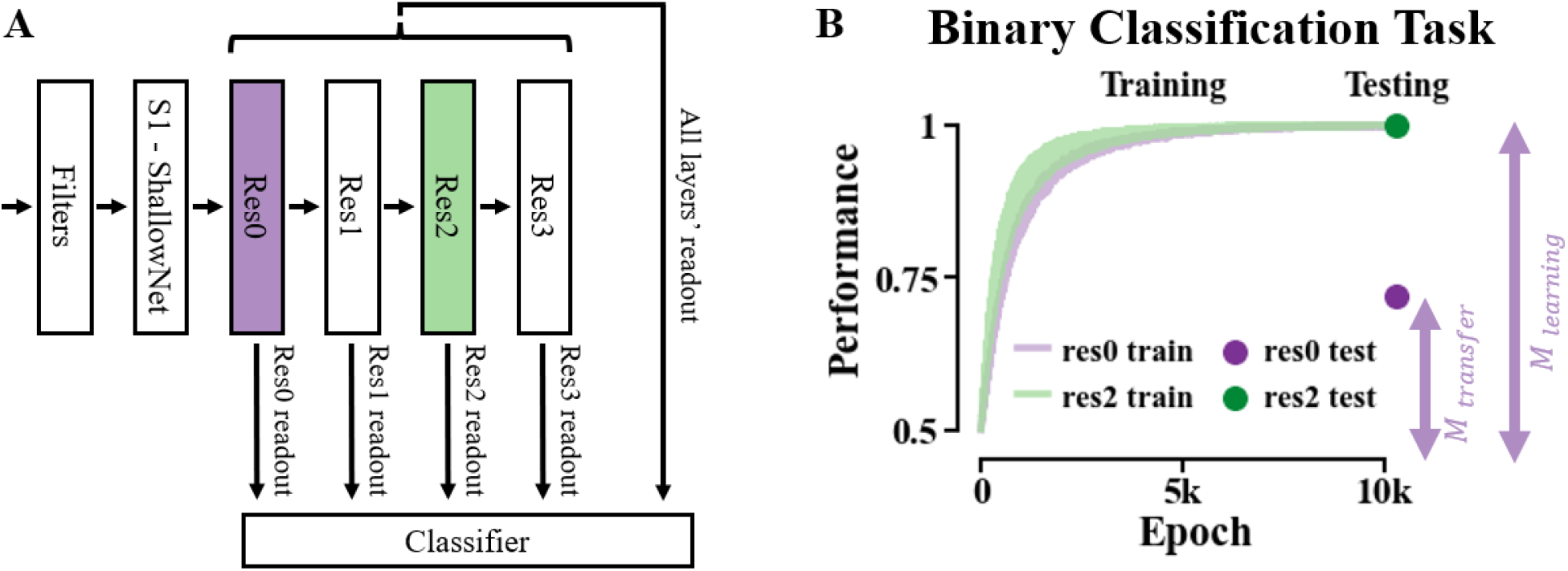
ANN readout modeling experiment. (A) Demonstration of the different readout mechanisms tested, using readouts from layers res0 through res3 individually, as well as all four layers concatenated together. (B) Example result showing classifier performance over 10,000 training epochs for the res3 readout, with performance on the location-transfer test shown at the end of training. *M*learning and *M*transfer for res0 stimuli are indicated in the plot (Eq. 6).

Across the six readout models, the specificity indices most closely resembled the human pattern when the classifier read out from deeper layers (res2 and res3) regardless of the training stimulus, rather than from early layers (res0, res1), from all layers combined, or from the stimulus-driven architecture (Fig 7). This qualitative correspondence between readout depth and the human behavioral pattern is consistent with the interpretation above: the specificity profile observed in human observers is best approximated by a model in which perceptual decisions are read out from higher-level stages. The other readout strategies led to specificity patterns after training that did not align with the human data. For example, unlike in human observers, the stimulus-driven readout produced equally minimal location specificity for res0 and res2 stimuli. Similarly, the res0 readout produced low location specificity for res0 stimuli and low stimulus-category specificity for res2 stimuli.

**Fig 7.**
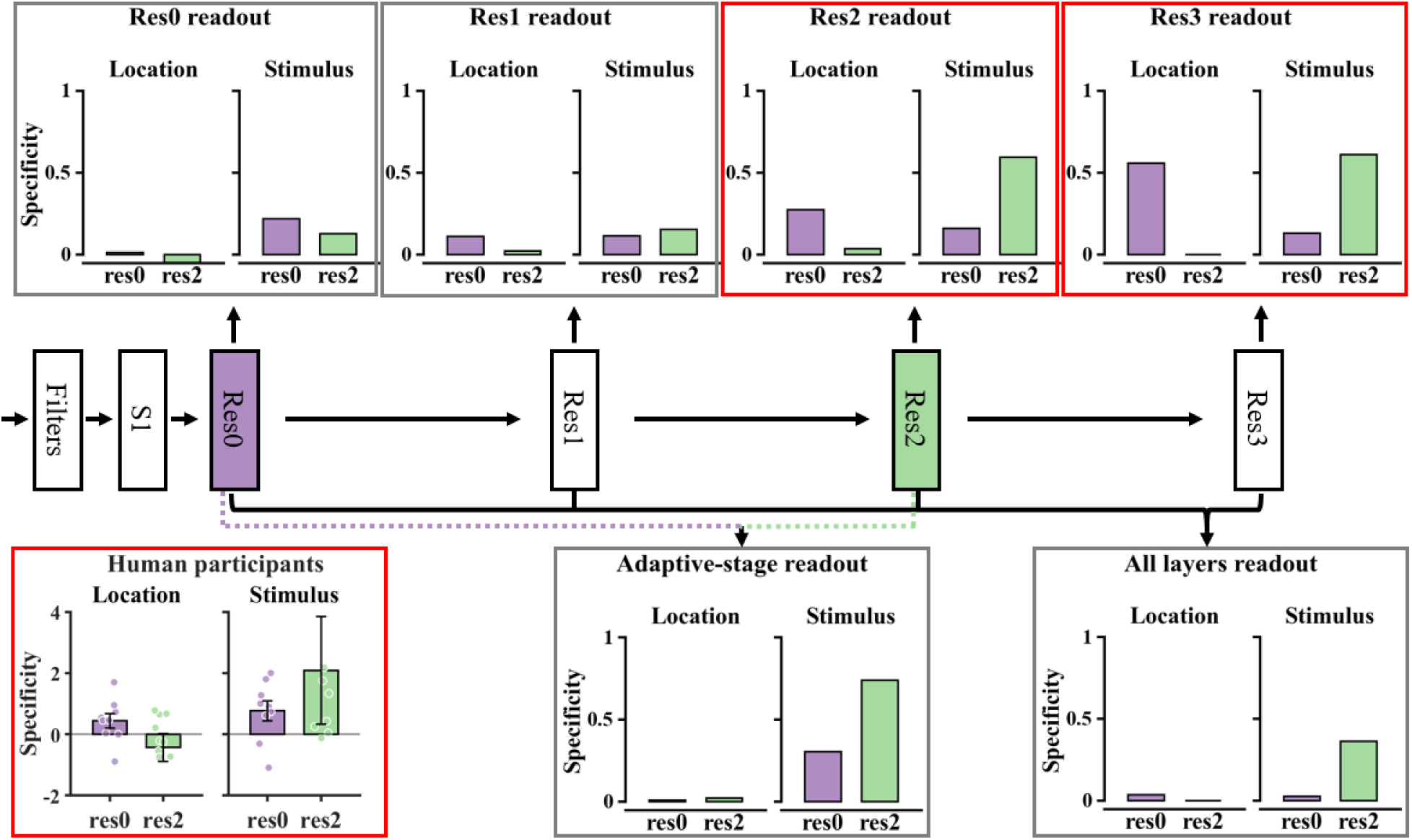
Specificity indices of ANN readout architectures compared with human observers. A schematic of DorsalNet shows the six readout architectures. Arrows indicate where each binary classifier read from: a single residual layer (res0, res1, res2 or res3), the combination of all layers (“All”), or the adaptive-stage architecture. For each readout architecture, classifiers were trained for 10,000 epochs, and specificity indices for simulated location and stimulus-category transfer were computed with the same procedure as for the human data (Section 2.5.2). Human specificity indices are shown alongside the model results. As the readout moves to later layers, the pattern of specificity indices becomes progressively more similar to the human pattern. The res2 and res3 readout models resemble the human data most closely; red outlines mark these two models and the human results. This correspondence is qualitative and based on visual inspection, not on a formal statistical test of model–human similarity. Participants’ data points falling outside the plotted axis range are omitted for figure readability.

## 4 Discussion

Our results suggest that the reweighting of readout during VPL operates at a high stage of the visual hierarchy, as the late-stage reweighting hypothesis proposes. Training improved performance for both stimulus types, but through different mechanisms. For res2 stimuli, which were tuned to a late stage, performance improved through a gain in sensitivity that transferred across retinal location. For res0 stimuli, which were tuned to an early stage, performance improved through a shift in response criterion that did not survive the transfer test. The ANN readout-architecture comparison converged on the same conclusion: the human specificity pattern was reproduced most closely by readouts from deep layers. Adaptive-stage reweighting, in which reweighting occurs at whichever stage is most informative about the trained stimulus, predicts sensitivity gains for both stimulus type. This prediction is difficult to reconcile with the human data, in which sensitivity improved only for res2 stimuli. The stimulus-driven readout model also failed to reproduce the human specificity pattern. In the terms introduced above, perceptual learning in this task resembles the lazy regime, in which a fixed late-stage representation is read out, and learning is confined to the readout weights. The sections below examine the sensitivity result, the criterion result, and the opposite specificity orderings in turn.

### 4.1 Sensitivity: learning is strongest, and most transferable, when the stimulus matches a higher-level readout

A default late-stage readout makes a prediction about performance before any learning: stimuli well separated at the stage being read out should be easier from the outset. Our data are in line with this prediction. Already in the first training session, res2-trained observers showed numerically higher sensitivity than res0-trained observers (Section 3.3). Reverse hierarchy theory makes the same prediction (Ahissar & Hochstein, 2004), a link we discuss further in Section 4.2. Human object recognition shows a similar dependence: its pattern of accuracy across tasks follows a weighted readout of a late stage of the object-processing hierarchy rather than of an intermediate stage, even though intermediate-stage populations outperform humans on some of those tasks (Majaj et al., 2015).

Training widened this early advantage. Res2 training produced large sensitivity gains, whereas res0 training produced only weak, non-significant gains. At transfer, feedback was removed, which does not necessarily abolish learning (Petrov et al., 2006; Seitz & Watanabe, 2003), and the res2 advantage remained significant at the untrained retinal location. Within the reweighting framework, learning adjusts the readout weights from a stable sensory representation to a decision unit (Dosher et al., 2013; Dosher & Lu, 1999; Petrov et al., 2005). Our results suggest that this readout draws on a late stage of the modeled pathway. Res2 pairs are well separated at this stage (Fig 1I); many of its units respond differently to the two stimuli of a pair, giving reweighting more informative units to draw on. Res0 pairs are poorly separated there (Fig 1H), leaving little difference for reweighting to emphasize. Learning was therefore strongest, and most transferable, when the trained stimuli were well separated at the stage being read out.

This dependence on separation at the read-out stage is consistent with a principle of learning from population codes: a readout learns fastest, and generalizes best, when the task aligns with the dominant dimensions of the code it reads (Bordelon & Pehlevan, 2022). Res2 stimuli were synthesized to be maximally distinct at res2, which makes their discrimination a task aligned with the late-stage code. A readout drawing on this stage should therefore learn res2 stimuli quickly and generalize what it learns, which is the pattern we observed.

This result also bears on the design of training curricula. Easy-to-hard curricula improve generalization in both humans and ANNs, because early training on easy conditions establishes a low-dimensional readout that later, harder training reuses (Volk et al., 2026). In our task, stimuli matched to late processing stages produced the largest and most transferable sensitivity gains. Such stimuli could therefore open a training curriculum, establishing a readout that later training on less well-matched stimuli can reuse.

### 4.2 Criterion carries learning when late-stage signals are weak

Signal detection theory separates performance into two quantities: sensitivity (d′), which reflects how well the two stimuli are separated in the observer’s internal decision variable, and response criterion (c), which reflects the observer’s tendency to favor one response over the other (Hautus et al., 2021) (Section 2.5.1). Most work on VPL specificity, including the reweighting accounts above, characterizes learning almost entirely in terms of sensitivity. Models of perceptual learning usually treat criterion as a fixed or unbiased quantity (Dosher & Lu, 1999; Gold et al., 1999; Petrov et al., 2005), and criterion is rarely analyzed as a dependent measure in its own right. A smaller body of work, however, indicates that criterion changes systematically with training and does not always change together with sensitivity (Aberg & Herzog, 2012; Jones et al., 2015; Wenger & Rasche, 2006). VPL may therefore involve more than changes in perceptual sensitivity, and criterion shifts can arise when the level of cortex engaged by a task extends beyond early sensory representations (Wenger et al., 2008).

Our res0 results show how criterion can carry learning when sensitivity cannot. At the start of training, observers in both categories were biased toward reporting "same" (Fig 5B). Our design presented equal numbers of same and different trials. Under this design, this bias lowers accuracy, and proportion correct reaches its maximum only when the criterion is unbiased (Hautus et al., 2021). For res0 stimuli, the late-stage readout receives only weakly separated signals (Fig 1H), leaving little room for reweighting to increase sensitivity. With this limited information available, moving the criterion toward the unbiased point becomes the main remaining way to improve accuracy. Res0-trained observers showed this pattern: their "same" bias declined significantly across training, and their performance improved without a significant sensitivity gain. Res2-trained observers, whose sensitivity increased, showed little change in bias.

The direction of this criterion shift agrees with previous reports. Supervised discrimination training with trial-by-trial feedback reduces a pre-existing response bias in naive observers in other perceptual domains (Jones et al., 2015), consistent with our design’s use of trial-by-trial feedback throughout training. The literature on criterion change in perceptual learning is not uniform, however: unsupervised training has instead been reported to shift criterion toward more liberal values (Wenger et al., 2008; Wenger & Rasche, 2006). The direction of criterion change may therefore depend on the presence and structure of feedback, rather than reflect a single fixed effect of practice.

The res0 bias reduction also did not survive the location-transfer test, in which two factors changed at once. Criterion shifts induced by biased feedback are specific to the trained position and orientation (Herzog et al., 2006), so a location change alone could undo a criterion adjustment. Criterion also depends on feedback: criteria track external feedback (Herzog & Fahle, 1999), and observers trained without feedback show higher decision bias than those trained with it, even when their sensitivity is matched (Petrov et al., 2006). Our design removed feedback at the same time as it changed location, so the reversal cannot be attributed to either factor alone. In either case, the res0 criterion adjustment proved less durable than the res2 sensitivity gain, which persisted despite both changes.

The criterion-based improvement with res0 training also bears on reverse hierarchy theory, an alternative account of learning-related shifts toward higher-level processing. Reverse hierarchy theory locates plasticity not in the weights linking a stable representation to a decision unit but in the representation itself: plasticity begins at high-level, abstract stages of processing and progresses to earlier, more fine-grained stages only as task demands require (Ahissar & Hochstein, 2004; Hochstein & Ahissar, 2002). This theory predicts that res0 training should eventually produce a sensitivity gain, but that this gain should take longer to appear. Across our three days of training, res0 performance did improve, yet sensitivity showed no significant gain; the improvement came instead from a shift in criterion. The early-stage plasticity that reverse hierarchy theory proposes may therefore still emerge with longer training. However, within the training period we tested, observers improved on res0 by adjusting their decision criterion, a pattern that reverse hierarchy theory alone cannot fully explain.

This argument, however, rests on an assumption that does not always hold. A change confined to criterion does not necessarily imply the absence of a perceptual change, since perceptual effects can in some cases manifest primarily as a criterion shift rather than a change in d′ (Witt et al., 2015). A purely criterion-based result therefore cannot rule out some degree of perceptual change; it shows only that any such change was too small to produce a detectable sensitivity gain. We therefore interpret the res0 pattern as a small, non-significant sensitivity change accompanied by a reliable criterion adjustment, rather than as either the complete absence of perceptual learning or strong evidence against reverse hierarchy theory.

### 4.3 Reconciling the opposite specificity orderings across transfer type

At first glance, res0 training was more specific to retinal location, whereas res2 training was more specific to stimulus category, and these orderings might seem to reflect two unrelated properties of the two conditions. We suggest instead that both follow from a single account in which readout is reweighted preferentially at a late stage of the pathway. Receptive fields grow from V1 through MT to MST (Section 1), and late-stage units therefore respond to motion over larger regions of the visual field and with more position-tolerant responses than early-stage units. Res2 stimuli target this late stage, so a sensitivity gain obtained by reweighting its readout should transfer across retinal location, as res2 sensitivity did. Res0 stimuli, by contrast, are poorly separated at this stage and produced little sensitivity gain, leaving little for a location change to disrupt. Their apparent location specificity therefore more plausibly reflects the reversion of the criterion adjustment described above than the loss of a location-specific sensitivity gain. As noted above, this reversion coincided with the removal of trial-by-trial feedback, so it may reflect the absence of feedback as well as the location change.

Stimulus-category transfer shows the reverse ordering, and the same late-stage readout accounts for it. Late-stage units integrate motion over wide regions and tolerate variation in its local structure, whereas res0 pairs were synthesized to differ chiefly in local structure, at an early stage. Res0 pairs are therefore poorly separated in late-stage activation space (Fig 1H), and readout weights adjusted during res2 training have little difference to act on once res0 stimuli replace res2 stimuli. Switching to res0 stimuli did not significantly reduce sensitivity, but it produced a marginal increase in response bias and the higher stimulus-category specificity index observed for res2, suggesting some cost of this switch. Res0 training, having produced little sensitivity gain to begin with, showed correspondingly little disruption of sensitivity when the stimulus category was switched. This account thus predicts the reversal in specificity ordering we observed across the two transfer types. It is also consistent with the ANN comparison, in which readouts from deep layers (res2 and res3), rather than from early layers, reproduced the human pattern of specificity indices most closely. This comparison was qualitative rather than a formal quantitative fit, and we therefore treat it as converging, rather than conclusive, evidence.

### 4.4 Late-stage readout in light of primate electrophysiology

Electrophysiological studies of perceptual learning in monkeys support readout reweighting and bear directly on which stage of the hierarchy is read out. In the dorsal pathway, training on random-dot direction discrimination changed motion-driven responses in the decision-related lateral intraparietal area (LIP) but not in MT, consistent with an increasingly selective readout of the most sensitive MT neurons (Law & Gold, 2008). This result is often cited in support of adaptive-stage reweighting, since MT represents random-dot motion well. MST neurons, however, can be as sensitive to random-dot motion as the animal’s behavior (Celebrini & Newsome, 1994), and changes in MST could not be ruled out (Law & Gold, 2008). For random-dot stimuli, readout from MT and late-stage readout from MST therefore make the same predictions; separating them requires stimuli that are informative at only one stage, as in the present study, or perturbation of MST.

The ventral pathway offers a parallel test. This pathway runs from V1 through V2 and V4 to posterior and anterior inferior temporal (IT) cortex (Kobatake & Tanaka, 1994). Along it, neurons respond to oriented edges in V1 (Hubel & Wiesel, 1968), to angles formed by pairs of contours in V2 (Ito & Komatsu, 2004), to boundary curvature at specific positions within a shape in V4 (Pasupathy & Connor, 2001), and to moderately complex object features in IT (Kobatake & Tanaka, 1994). Receptive fields grow along the same progression (Kobatake & Tanaka, 1994). After extensive training on noisy oriented gratings, a stimulus classically associated with early stages, neurons in posterior IT (PIT) discriminated the trained orientations better than untrained ones (Adab et al., 2014), and perturbing or inactivating PIT impaired the discrimination (Adab & Vogels, 2016). A late stage thus contributes causally to decisions about even low-level stimuli, as late-stage reweighting predicts. The small effect of PIT inactivation indicates that other areas contribute as well (Adab & Vogels, 2016), consistent with a readout weighted toward late stages rather than confined to them.

The strongest challenge to a default late-stage readout comes from inactivation of MT. After months of training on drifting gratings, MT inactivation barely affected grating discrimination; after training with random dots, the same inactivation strongly impaired it, and, in one animal, renewed grating training gradually reversed this effect (Liu & Pack, 2017). Training can therefore move the readout between MT and lower-level areas, as adaptive-stage reweighting proposes. These findings nonetheless fit a late-stage default within reverse hierarchy theory, in which decisions are read out by default from higher areas such as MT or MST and shift to lower areas only after extended training on difficult discriminations (Ahissar & Hochstein, 2004; Liu & Pack, 2017). The monkeys trained for months before testing, whereas our observers trained for three days, a duration that may be too short to move the readout to an early stage and that is consistent with the weak res0 sensitivity gain we observed. Moreover, the lower-level readout was inferred from the absence of an inactivation effect and from behavioral measures of spatial integration, rather than from perturbing early areas or MST, so grating signals may also have reached later stages through routes that bypass the inactivated region of MT. Together, these studies are consistent with a readout that defaults to late stages and that prolonged training can shift toward earlier ones.

### 4.5 Late-stage reweighting as a lazy, energy-efficient form of learning

In the terminology of learning dynamics in ANNs, late-stage reweighting parallels the lazy regime, in which internal representations remain essentially fixed, most network parameters hardly change, and learning is confined largely to the readout weights (Chizat et al., 2019; Farrell et al., 2023; Flesch et al., 2022). Adaptive-stage reweighting, by contrast, relocates reweighting to whichever stage shows the most distinct activations for each trained stimulus, and therefore requires reweighting readout weights at many levels of the hierarchy. Our results favor this lazy-like form of learning. Reweighting remained at a late stage even when the trained stimuli were most distinct at an early stage, and observers compensated for the resulting lack of sensitivity by adjusting their criterion rather than by relocating reweighting to res0.

A lazy regime may confer an energetic advantage. Plasticity modifies fewer synapses when it is confined to the readout weights of a single late stage than when it is distributed across the hierarchy, and synaptic plasticity is metabolically costly (Li & van Rossum, 2020; van Rossum & Pache, 2024). In Drosophila, forming long-term memory reduces survival (Mery & Kawecki, 2005), and starved flies suppress the formation of this costly memory (Plaçais & Preat, 2013). More broadly, neural signaling is energetically expensive (Attwell & Laughlin, 2001), energy limitation has acted as a selective pressure on the evolution of sensory systems (Niven & Laughlin, 2008), and mouse visual cortex reduces its coding precision to save energy during food scarcity (Padamsey et al., 2022). Evolution may therefore have favored learning mechanisms that achieve behavioral gains with as few synaptic changes as possible (van Rossum & Pache, 2024), and reweighting readout from a fixed late stage is one such mechanism.

This lazy form of learning may characterize only the early phase of learning. In ANNs, networks can remain in the lazy regime early in training and depart from it only later (Kumar et al., 2024), and our three days of training may capture only this early phase. Whether longer training shifts reweighting to earlier stages of the hierarchy, and whether the lazy regime reflects an energetic constraint, remain open questions. We did not measure energy expenditure, and this interpretation remains speculative.

### 4.6 Limitations

Two limitations qualify these conclusions. First, the stage each stimulus category targets was inferred from DorsalNet, whose layers align with successive areas of the primate dorsal pathway (Mineault et al., 2021). On this basis, res0 stimuli are expected to engage early areas such as V1/MT, and res2 stimuli later areas such as MST. Confirming this correspondence directly will require electrophysiological recordings in primates presented with these stimuli. Second, res0- and res2-trained observers formed separate, non-overlapping groups, so any difference between the two conditions could partly reflect pre-existing individual differences between groups (e.g., in baseline motion sensitivity or task engagement) rather than the stimulus manipulation alone. A within-subject design, in which each observer is trained on both stimulus categories with order counterbalanced across participants, would strengthen the causal interpretation of these effects. Such a design, however, introduces its own complications, including order and learning-to-learn effects and a substantially longer testing protocol.

## 5 Conclusion

This study used model-based stimulus synthesis using DorsalNet to ask at which stage of the visual hierarchy the readout supporting perceptual decisions is reweighted during visual perceptual learning. Training improved performance for both stimulus categories, but through different mechanisms. Training with stimuli synthesized to be maximally discernible by a late network stage (res2) significantly increased sensitivity, and this gain persisted at a new retinal location without trial-by-trial feedback. Training with stimuli synthesized to be maximally discernible by an early stage (res0) produced only a weak, non-significant change in sensitivity but a significant reduction in response bias; this reduction reverted at the new retinal location, where feedback was also withheld. Specificity indices for location and stimulus-category transfer showed opposite orderings for the two categories, and ANN classifiers qualitatively reproduced this pattern most closely when their readout was drawn from late, rather than early, layers. Together, these findings suggest that readout in this task is reweighted at a comparatively late stage of the dorsal pathway, whatever the trained stimulus. How well the trained stimulus matches this stage determines how learning is expressed: well-matched stimuli yield sensitivity gains, whereas poorly matched stimuli yield only weak discrimination, which observers exploit by shifting their decision criterion. Sensitivity and criterion therefore reflect at least partially distinct learning mechanisms rather than interchangeable indices of a single process.

## 6 Supplementary materials

**S1 Appendix.** The synthesized stimuli, together with the code and documentation for stimulus synthesis and ANN training, are available at https://younesvali.com/projects/dorsalnet/. If the link is unavailable, go to younesvali.com, open the Projects tab, and select “Hierarchical control of visual perceptual learning with model-synthesized stimuli”.

## 7 Acknowledgment

We thank Maria Haddad and Patrick Mineault for their assistance in setting up the initial phase of stimulus generation and Yasaman Bagi for her assistance in setting up the initial phase of data collection. This work was supported by NSERC Discovery Grants (RGPIN-2023-03875 to SB; RGPIN/04333-2017) to CCP, an IVADO Exploratory Project grant (Explo24CO-3750823649) to SB and CCP, and an Excellence Scholarship from UNIQUE (Unifying Neuroscience and Artificial Intelligence - Québec) awarded to YV. We also thank all participants for their time.

## References

1. Aberg, K. C., & Herzog, M. H. (2012). Different types of feedback change decision criterion and sensitivity differently in perceptual learning. Journal of Vision, 12(3), 3–3.

2. Adab, H. Z., Popivanov, I. D., Vanduffel, W., & Vogels, R. (2014). Perceptual Learning of Simple Stimuli Modifies Stimulus Representations in Posterior Inferior Temporal Cortex. Journal of Cognitive Neuroscience, 26(10), 2187–2200. 10.1162/jocn_a_00641

3. Adab, H. Z., & Vogels, R. (2016). Perturbation of Posterior Inferior Temporal Cortical Activity Impairs Coarse Orientation Discrimination. Cerebral Cortex, 26(9), 3814–3827. 10.1093/cercor/bhv178

4. Ahissar, M., & Hochstein, S. (2004). The reverse hierarchy theory of visual perceptual learning. Trends in Cognitive Sciences, 8(10), 457–464. 10.1016/j.tics.2004.08.011

5. Albright, T. D., & Desimone, R. (1987). Local precision of visuotopic organization in the middle temporal area (MT) of the macaque. Experimental Brain Research, 65(3), 582–592. 10.1007/BF00235981

6. Attwell, D., & Laughlin, S. B. (2001). An Energy Budget for Signaling in the Grey Matter of the Brain. Journal of Cerebral Blood Flow & Metabolism, 21(10), 1133–1145. 10.1097/00004647-200110000-00001

7. Bakhtiari, S., Awada, A., & Pack, C. C. (2020). Influence of stimulus complexity on the specificity of visual perceptual learning. Journal of Vision, 20(6), 13. 10.1167/jov.20.6.13

8. Bakhtiari, S., Mineault, P., Lillicrap, T., Pack, C., & Richards, B. (2021). The functional specialization of visual cortex emerges from training parallel pathways with self-supervised predictive learning. Advances in Neural Information Processing Systems, 34, 25164–25178.

9. Ball, K., & Sekuler, R. (1982). A Specific and Enduring Improvement in Visual Motion Discrimination. *Science (New York*, N.Y.), 218(4573), 697–698. doi:10.1126/science.7134968

10. Bashivan, P., Kar, K., & DiCarlo, J. J. (2019). Neural population control via deep image synthesis. *Science (New York*, N.Y.), 364(6439). 10.1126/science.aav9436

11. Bordelon, B., & Pehlevan, C. (2022). Population codes enable learning from few examples by shaping inductive bias. eLife, 11, e78606. 10.7554/eLife.78606

12. Celebrini, S., & Newsome, W. T. (1994). Neuronal and psychophysical sensitivity to motion signals in extrastriate area MST of the macaque monkey. The Journal of Neuroscience, 14(7), 4109–4124.

13. Cheng, Y.-A., Sanayei, M., Chen, X., Jia, K., Li, S., Fang, F., Watanabe, T., Thiele, A., & Zhang, R.-Y. (2025). A neural geometry approach comprehensively explains apparently conflicting models of visual perceptual learning. Nature Human Behaviour, 9(5), 1023–1040. 10.1038/s41562-025-02149-x

14. Chizat, L., Oyallon, E., & Bach, F. (2019). On lazy training in differentiable programming. Advances in Neural Information Processing Systems,

15. Cohen, G., & Weinshall, D. (2017). Hidden layers in perceptual learning. Proceedings of the IEEE Conference on Computer Vision and Pattern Recognition,

16. Desimone, R., & Ungerleider, L. G. (1986). Multiple visual areas in the caudal superior temporal sulcus of the macaque. Journal of Comparative Neurology, 248(2), 164–189.

17. DiCarlo, J. J., Zoccolan, D., & Rust, N. C. (2012). How does the brain solve visual object recognition? Neuron, 73(3), 415–434.

18. Dosher, B., & Lu, Z.-L. (2020). Perceptual learning: How experience shapes visual perception. MIT Press.

19. Dosher, B. A., Jeter, P., Liu, J., & Lu, Z.-L. (2013). An integrated reweighting theory of perceptual learning. Proceedings of the National Academy of Sciences, 110(33), 13678–13683.

20. Dosher, B. A., & Lu, Z.-L. (1999). Mechanisms of perceptual learning. Vision Research, 39(19), 3197–3221. 10.1016/S0042-6989(99)00059-0

21. Duffy, C. J., & Wurtz, R. H. (1991). Sensitivity of MST neurons to optic flow stimuli. I. A continuum of response selectivity to large-field stimuli. Journal of Neurophysiology, 65(6), 1329–1345. 10.1152/jn.1991.65.6.1329

22. Farrell, M., Recanatesi, S., & Shea-Brown, E. (2023). From lazy to rich to exclusive task representations in neural networks and neural codes. Current Opinion in Neurobiology, 83, 102780. 10.1016/j.conb.2023.102780

23. Fiorentini, A., & Berardi, N. (1980). Perceptual learning specific for orientation and spatial frequency. Nature, 287(5777), 43–44. 10.1038/287043a0

24. Flesch, T., Juechems, K., Dumbalska, T., Saxe, A., & Summerfield, C. (2022). Orthogonal representations for robust context-dependent task performance in brains and neural networks. Neuron, 110(7), 1258–1270.e1211. 10.1016/j.neuron.2022.01.005

25. Geirhos, R., Narayanappa, K., Mitzkus, B., Thieringer, T., Bethge, M., Wichmann, F. A., & Brendel, W. (2021). Partial success in closing the gap between human and machine vision. Advances in Neural Information Processing Systems, 34, 23885–23899.

26. Gold, J., Bennett, P. J., & Sekuler, A. B. (1999). Signal but not noise changes with perceptual learning. Nature, 402(6758), 176–178. 10.1038/46027

27. Hautus, M. J., Macmillan, N. A., & Creelman, C. D. (2021). Detection theory: A user’s guide (3rd ed.). Routledge.

28. Herzog, M. H., Ewald, K. R. F., Hermens, F., & Fahle, M. (2006). Reverse feedback induces position and orientation specific changes. Vision Research, 46(22), 3761–3770. 10.1016/j.visres.2006.04.024

29. Herzog, M. H., & Fahle, M. (1999). Effects of biased feedback on learning and deciding in a vernier discrimination task. Vision Research, 39(25), 4232–4243. 10.1016/S0042-6989(99)00138-8

30. Hochstein, S., & Ahissar, M. (2002). View from the top: Hierarchies and reverse hierarchies in the visual system. Neuron, 36(5), 791–804.

31. Hubel, D. H., & Wiesel, T. N. (1968). Receptive fields and functional architecture of monkey striate cortex. The Journal of Physiology, 195(1), 215–243.

32. Ito, M., & Komatsu, H. (2004). Representation of Angles Embedded within Contour Stimuli in Area V2 of Macaque Monkeys. Journal of Neuroscience, 24(13), 3313–3324. 10.1523/JNEUROSCI.4364-03.2004

33. Jones, P. R., Moore, D. R., Shub, D. E., & Amitay, S. (2015). The role of response bias in perceptual learning. *Journal of Experimental Psychology: Learning*, Memory, and Cognition, 41(5), 1456.

34. Khaligh-Razavi, S.-M., & Kriegeskorte, N. (2014). Deep Supervised, but Not Unsupervised, Models May Explain IT Cortical Representation. PLOS Computational Biology, 10(11), e1003915. 10.1371/journal.pcbi.1003915

35. Khawaja, F. A., Liu, L. D., & Pack, C. C. (2013). Responses of MST neurons to plaid stimuli. Journal of Neurophysiology, 110(1), 63–74.

36. Kingma, D. P., & Ba, J. (2014). Adam: A method for stochastic optimization. arXiv preprint arXiv:1412.6980.

37. Kobatake, E., & Tanaka, K. (1994). Neuronal selectivities to complex object features in the ventral visual pathway of the macaque cerebral cortex. Journal of Neurophysiology, 71(3), 856–867. 10.1152/jn.1994.71.3.856

38. Kumar, T., Bordelon, B., Gershman, S., & Pehlevan, C. (2024). Grokking as the transition from lazy to rich training dynamics. International Conference on Learning Representations,

39. Laamerad, P., Awada, A., Pack, C. C., & Bakhtiari, S. (2024). Asymmetric stimulus representations bias visual perceptual learning. Journal of Vision, 24(1), 10–10.

40. Laamerad, P., Krause, M. R., Guitton, D., & Pack, C. C. (2025). Inactivation of primate cortex reveals inductive biases in visual learning. Curr Biol, 35(19), 4699–4713.e4696. 10.1016/j.cub.2025.08.027

41. Law, C.-T., & Gold, J. I. (2008). Neural correlates of perceptual learning in a sensory-motor, but not a sensory, cortical area. Nature Neuroscience, 11(4), 505–513.

42. Li, H. L., & van Rossum, M. C. W. (2020). Energy efficient synaptic plasticity. eLife, 9, e50804. 10.7554/eLife.50804

43. Liu, L. D., & Pack, C. C. (2017). The Contribution of Area MT to Visual Motion Perception Depends on Training. Neuron, 95(2), 436–446.e433. 10.1016/j.neuron.2017.06.024

44. Liu, X., Lyu, M., Yu, C., & Wu, S. (2024). To learn or not to learn, that is the question—a feature-task dual learning model of perceptual learning. Advances in Neural Information Processing Systems, 37, 37736–37764.

45. Lu, Z.-L., & Dosher, B. A. (2022). Current directions in visual perceptual learning. Nature Reviews Psychology, 1(11), 654–668. 10.1038/s44159-022-00107-2

46. Majaj, N. J., Hong, H., Solomon, E. A., & DiCarlo, J. J. (2015). Simple learned weighted sums of inferior temporal neuronal firing rates accurately predict human core object recognition performance. The Journal of Neuroscience, 35(39), 13402–13418.

47. Manenti, G. L., Dizaji, A. S., & Schwiedrzik, C. M. (2023). Variability in training unlocks generalization in visual perceptual learning through invariant representations. Current Biology, 33(5), 817–826.e813. 10.1016/j.cub.2023.01.011

48. Maunsell, J. H., & Essen, D. C. v. (1983). The connections of the middle temporal visual area (MT) and their relationship to a cortical hierarchy in the macaque monkey. Journal of Neuroscience, 3(12), 2563–2586. 10.1523/JNEUROSCI.03-12-02563.1983

49. Mery, F., & Kawecki, T. J. (2005). A Cost of Long-Term Memory in Drosophila. *Science (New York*, N.Y.), 308(5725), 1148–1148. doi:10.1126/science.1111331

50. Mineault, P., Bakhtiari, S., Richards, B., & Pack, C. (2021, 2021). Your head is there to move you around: Goal-driven models of the primate dorsal pathway.

51. Mineault, P. J., Khawaja, F. A., Butts, D. A., & Pack, C. C. (2012). Hierarchical processing of complex motion along the primate dorsal visual pathway. Proceedings of the National Academy of Sciences, 109(16), E972–E980. 10.1073/pnas.1115685109

52. Niven, J. E., & Laughlin, S. B. (2008). Energy limitation as a selective pressure on the evolution of sensory systems. Journal of Experimental Biology, 211(11), 1792–1804.

53. Olah, C., Mordvintsev, A., & Schubert, L. (2017). Feature Visualization. Distill, 2(11), e7. 10.23915/distill.00007

54. Padamsey, Z., Katsanevaki, D., Dupuy, N., & Rochefort, N. L. (2022). Neocortex saves energy by reducing coding precision during food scarcity. Neuron, 110(2), 280–296.e210. 10.1016/j.neuron.2021.10.024

55. Pasupathy, A., & Connor, C. E. (2001). Shape Representation in Area V4: Position-Specific Tuning for Boundary Conformation. Journal of Neurophysiology, 86(5), 2505–2519. 10.1152/jn.2001.86.5.2505

56. Petrov, A. A., Dosher, B. A., & Lu, Z.-L. (2005). The dynamics of perceptual learning: an incremental reweighting model. Psychological review, 112(4), 715.

57. Petrov, A. A., Dosher, B. A., & Lu, Z.-L. (2006). Perceptual learning without feedback in non-stationary contexts: Data and model. Vision Research, 46(19), 3177–3197. 10.1016/j.visres.2006.03.022

58. Plaçais, P.-Y., & Preat, T. (2013). To Favor Survival Under Food Shortage, the Brain Disables Costly Memory. *Science (New York*, N.Y.), 339(6118), 440–442. doi:10.1126/science.1226018

59. Ponce, C. R., Xiao, W., Schade, P. F., Hartmann, T. S., Kreiman, G., & Livingstone, M. S. (2019). Evolving Images for Visual Neurons Using a Deep Generative Network Reveals Coding Principles and Neuronal Preferences. Cell, 177(4), 999–1009.e1010. 10.1016/j.cell.2019.04.005

60. Saito, H., Yukie, M., Tanaka, K., Hikosaka, K., Fukada, Y., & Iwai, E. (1986). Integration of direction signals of image motion in the superior temporal sulcus of the macaque monkey. Journal of Neuroscience, 6(1), 145–157. 10.1523/JNEUROSCI.06-01-00145.1986

61. Schrimpf, M., Kubilius, J., Hong, H., Majaj, N. J., Rajalingham, R., Issa, E. B., Kar, K., Bashivan, P., Prescott-Roy, J., Geiger, F., Schmidt, K., Yamins, D. L. K., & DiCarlo, J. J. (2020). Brain-Score: Which Artificial Neural Network for Object Recognition is most Brain-Like? https://www.biorxiv.org/content/10.1101/407007v2

62. Schrimpf, M., Kubilius, J., Lee, M. J., Ratan Murty, N. A., Ajemian, R., & DiCarlo, J. J. (2020). Integrative Benchmarking to Advance Neurally Mechanistic Models of Human Intelligence. Neuron, 108(3), 413–423. 10.1016/j.neuron.2020.07.040

63. Seitz, A. R., & Watanabe, T. (2003). Is subliminal learning really passive? Nature, 422(6927), 36–36. 10.1038/422036a

64. Shah, S., Dey, D., Lovett, C., & Kapoor, A. (2017). Airsim: High-fidelity visual and physical simulation for autonomous vehicles. Field and service robotics: Results of the 11th international conference,

65. Shen, S., Sun, Y., Lu, J., Li, C., Chen, Q., Mo, C., Fang, F., & Zhang, X. (2024). Profiles of visual perceptual learning in feature space. iScience, 27(3), 109128. 10.1016/j.isci.2024.109128

66. Sotiropoulos, G., Seitz, A. R., & Seriès, P. (2011). Perceptual learning in visual hyperacuity: A reweighting model. Vision Research, 51(6), 585–599. 10.1016/j.visres.2011.02.004

67. Tanaka, K., Hikosaka, K., Saito, H., Yukie, M., Fukada, Y., & Iwai, E. (1986). Analysis of local and wide-field movements in the superior temporal visual areas of the macaque monkey. Journal of Neuroscience, 6(1), 134–144. 10.1523/JNEUROSCI.06-01-00134.1986

68. van Rossum, M. C., & Pache, A. (2024). Competitive plasticity to reduce the energetic costs of learning. PLOS Computational Biology, 20(10), e1012553.

69. Volk, C., Pack, C. C., & Bakhtiari, S. (2026). The curriculum effect in visual learning: the role of readout dimensionality. PLOS Computational Biology, 22(7), e1014553.

70. Walker, E. Y., Sinz, F. H., Cobos, E., Muhammad, T., Froudarakis, E., Fahey, P. G., Ecker, A. S., Reimer, J., Pitkow, X., & Tolias, A. S. (2019). Inception loops discover what excites neurons most using deep predictive models. Nature Neuroscience, 22(12), 2060–2065. 10.1038/s41593-019-0517-x

71. Wenger, M. J., Copeland, A. M., Bittner, J. L., & Thomas, R. D. (2008). Evidence for criterion shifts in visual perceptual learning: Data and implications. Perception & Psychophysics, 70(7), 1248–1273.

72. Wenger, M. J., & Rasche, C. (2006). Perceptual learning in contrast detection: Presence and cost of shifts in response criteria. Psychonomic bulletin & review, 13(4), 656–661.

73. Wenliang, L. K., & Seitz, A. R. (2018). Deep Neural Networks for Modeling Visual Perceptual Learning. The Journal of Neuroscience, 38(27), 6028–6044. 10.1523/jneurosci.1620-17.2018

74. Witt, J. K., Taylor, J. E. T., Sugovic, M., & Wixted, J. T. (2015). Signal detection measures cannot distinguish perceptual biases from response biases. Perception, 44(3), 289–300.

75. Yamins, D. L. K., Hong, H., Cadieu, C. F., Solomon, E. A., Seibert, D., & DiCarlo, J. J. (2014). Performance-optimized hierarchical models predict neural responses in higher visual cortex. Proceedings of the National Academy of Sciences, 111(23), 8619–8624. 10.1073/pnas.1403112111

